# Structural basis of stereochemical promiscuity by an umami taste receptor ortholog, Tas1r1/Tas1r3 from pufferfish

**DOI:** 10.1101/2025.11.24.689453

**Authors:** Rakuto Mizoguchi, Yasuka Toda, Mana Nagae, Takashi Yoshida, Hiroaki Matsuura, Kunio Hirata, Yohei Miyanoiri, Maiko Hosotani, Yuji Ashikawa, Chiaki Ito, Naotaka Tsutsumi, Norihisa Yasui, Yoshiro Ishimaru, Atsuko Yamashita

## Abstract

Taste receptor type 1 (TAS1R), which consists of sweet and umami receptors in humans, senses nutrients (such as sugars and amino acids) with substrate specificity that is often broad and varies among animals and subtypes. However, the structural basis for achieving diverse specificities remains largely elusive. Here, we present the crystal structure of the ligand-binding domain (LBD) of Tas1r1/Tas1r3 heterodimer from pufferfish, an ortholog of the human umami taste receptor. The overall structure of Tas1r1/Tas1r3LBD resembles previously reported TAS1R structures, indicating a conserved core architecture within the family. Nevertheless, pufferfish Tas1r1/Tas1r3 was found to bind and respond to both L- and D-amino acids, even though TAS1Rs are considered to respond to either enantiomer. Structural and mutational analyses revealed that this non-rigorous stereochemical recognition is attributed to inter-subdomain interactions that latch the cleft containing the amino acid-binding site. These interactions prevent the cleft from fully opening, thereby stabilizing the active conformation, even if the ligand shares a different chirality. These results suggest that different substrate specificities in TAS1Rs can be acquired not only by the gain or loss of direct interactions with the substrate but also by the gain of intramolecular interactions, which alter the conformational equilibrium of the receptor.

## Introduction

Taste senses chemicals in foods and is therefore intrinsically linked to the feeding behaviors of animals. Taste modalities detecting nutrients, such as sweet and umami (savory) taste sensing sugars and amino acids, are perceived as preferable tastes and promote their consumption, while those detecting potentially harmful substances, such as bitter and sour taste sensing chemicals in toxic substances and deteriorated foods, are perceived as unpreferable tastes and promote their avoidance.

Taste receptor type 1 (TAS1R or T1R) is a protein family responsible for nutrient detection and preferable taste sensation. In mammals such as humans, the receptors for two modalities, sweet and umami (savory taste), are composed of heterodimeric TAS1Rs; TAS1R1/TAS1R3 serves as the umami receptor recognizing L-amino acids and nucleotides, whereas TAS1R2/TAS1R3 serves as the sweet receptor recognizing sugars and D-amino acids ^1-3^. TAS1Rs are conserved among vertebrates, and at least those characterized to date commonly respond to sugars, amino acids, and/or nucleotides. A recent genome-wide analysis of TAS1R genes revealed that there are 11 TAS1R members in vertebrates, and each animal species possesses a limited set of TAS1Rs, as found in mammals, which share three members ^4^. Among them, TAS1R1 is the most commonly equipped TAS1R member across the bony vertebrates classes ^4,5^. Nevertheless, the substrate specificities of the TAS1R1/TAS1R3 receptors are diverse and distinct in different animal species. For example, TAS1R1/TAS1R3 in humans, as well as in other leaf-eating primates, show high responses to L-glutamate but weak responses to 5′-ribonucleotides, whereas those in insect-eating primates show high responses to 5′- ribonucleotides according with the contents of the chemicals in their foods ^6^. Among birds, which only have TAS1R1/TAS1R3 in the entire class, the receptors in insect-eating species, such as chimney swift, respond to amino acids, whereas those in nectar-eating hummingbirds respond to sugars in addition to amino acids ^7,8^. These results indicate that TAS1Rs in each animal species acquire the substrate specificity suitable for their food habits through molecular evolution. Another characteristic of TAS1Rs is that they often show broad substrate specificities; for example, although human TAS1R1/TAS1R3 shows specificity to L-glutamate and aspartate, TAS1R1/TAS1R3 in other animals (e.g. mice) responds to a wide range of L-amino acids ^2,3,9^.

TAS1Rs belong to the class C G protein-coupled receptor (GPCR) family, in which the ligand-binding domain (LBD) on the extracellular side upstream of the hepta-helical transmembrane region is responsible for agonist binding ^10^. Clues for understanding the substrate specificity of TAS1Rs were initially provided by the crystallographic structure of the LBD of Tas1r2a/Tas1r3 (T1r2a/T1r3) in medaka fish, which was the only TAS1R structure to be solved until recently ^11^. L-amino acids, substrates for medaka Tas1r2a/Tas1r3, were observed to bind at the substrate-binding sites located in the inter-subdomain clefts of both subunits ^11^, similar to the other class C GPCR structures. Recent accumulated structural information regarding the full-length class C GPCRs suggested an activation scheme of the receptors initiated by a conformational change in the LBD ^12-20^, in which agonist binding induces closure of the cavity between the two subdomains in the LBD, thus rearranging the downstream transmembrane region by adopting a suitable dimer configuration for cytosolic G-protein activation. Indeed, medaka Tas1r2a/Tas1r3LBD showed a conformational change upon binding of an amino acid ^11,21^, essentially in a manner similar to that observed in other class C GPCRs. Since medaka Tas1r2a/Tas1r3 binds and responds to a wide array of L-amino acids ^22,23^, the binding sites provide the residues that strictly recognize the common groups in amino acids as α-amino and carboxy groups together with the spaces amenable to accommodate a substituent group exhibiting various physicochemical properties ^11^. Recently, breakthrough studies for the elucidation of the full-length structures of the human sweet receptor TAS1R2/TAS1R3 were reported, which showed the binding of sweet substances, such as sucralose or advantame, at the same binding site where the amino acid ligands were observed in the medaka receptor ^24-26^. Nevertheless, since the currently available structural information of TAS1Rs is limited to these two receptors, the structural basis for the recognition of a variety of chemicals and their specificity in TAS1Rs remains largely unknown.

In this study, we determined the crystallographic structure of another member of the TAS1Rs, the LBD of pufferfish Tas1r1/Tas1r3, a heterodimer of TAS1R1 and TAS1R3B group proteins, in which the former subunit is an ortholog of mammalian umami taste receptors ^4^. We found that the broadness of the substrate specificity of this receptor is extended to stereochemistry: the receptor binds and responds to D-amino acids in addition to the L-enantiomers, unlike many other biological macromolecules, such as enzymes, showing rigorous stereospecificities. The structures of pufferfish Tas1r1/Tas1r3LBD bound to L- and D-amino acids, together with recombinant protein-based ligand binding analysis and cell signaling assays, pinpointed the specific inter-subdomain interactions stabilizing the active conformation, thus allowing signal transduction even when the bound ligand has a different chirality.

## Results

### Structure of the ligand-binding domain of pufferfish Tas1r1/Tas1r3

Structural analysis of TAS1R1/TAS1R3 was hampered because most receptor proteins from various species showed no noticeable recombinant expression in properly folded states during our expression screening ^27^. Nevertheless, we found that Tas1r1LBD and Tas1r3LBD from pufferfish, *Takifugu rubripes*, exhibit plausible expression with proper extracellular localization. We further introduced the Cys242S mutation in Tas1r1 and Cys238S mutation in Tas1r3 to eliminate free cysteine residues at the molecular surface to avoid unexpected intermolecular disulfide formation (Extended Data Fig. 1; see Methods), and this backbone was employed for further analyses using the LBD protein sample. The co-introduction of these two genes into *Drosophila* S2 cells and the establishment of a stable high-expression clone ^28^ enabled the preparation of heterodimeric protein samples for structural analysis (Extended Data Fig. 1). Pufferfish Tas1r1 shares an amino acid sequence identity with human TAS1R1 as 40%, while with medaka Tas1r2a and 30% and with human TAS1R2 of 35% (Extended Data Fig. 2), which is consistent with the notion that this protein is an ortholog of mammalian umami-receptor subunit TAS1R1 ^4,5,29^. The crystallographic structure of Tas1r1/Tas1r3LBD was solved in complex with L-alanine (Extended Data Table 1), an amino acid that induces taste nerve responses in pufferfish ^30,31^ and serves as a typical taste substance for TAS1R1/TAS1R3 in many animals ^2,7,32^.

The Tas1r1LBD and Tas1r3LBD subunits share the bi-lobal architecture referred to as the Venus-flytrap module, similar to medaka Tas1r2a/Tas1r3LBD ^11^ (a heterodimer of TAS1R2B/TAS1R3B member proteins; Cα rmsd: 1.6 Å), human TAS1R2/TAS1R3 ^24-26^ (a heterodimer of TAS1R2A/TAS1R3A member proteins; Cα rmsd: 2.8 Å), and the other class C GPCR LBDs ^10^ (Fig. 1A–C). The two subunits in the heterodimer are tethered to each other by an asymmetric intermolecular disulfide bond between Cys366 in Tas1r1 and Cys131 in Tas1r3 (Fig. 1D, Extended Data Fig. 1B, C), similar to those observed in medaka Tas1r2a/Tas1r3LBD and human TAS1R2/TAS1R3 heterodimers (Extended Data Fig. 3A, B). The two subunits adopt a compact dimerization manner with clefts between the two subdomains, LB1 and LB2, in both subunit closing, namely the “closed” state, which is considered as a conformation required for receptor activation ^10^. The conformation is similar to those of medaka Tas1r2a/Tas1r3 in complex with various amino acids serving as agonists for the receptor (Fig. 1B, E) and human TAS1R2 in complex with sucralose (Fig. 1C, F) or advantame, while human TAS1R3 adopted the “open” state due to the absence of sweet substance binding (Fig. 1F). L-alanine binding was observed at the inner end of the cleft between LB1 and LB2 in each subunit (Extended Data Fig. 4), which corresponds to the amino acid-binding site in medaka Tas1r2a/Tas1r3 (Fig. 1B, E), the sweet substance-binding site in human TAS1R2 (Fig. 1C, F), and orthosteric ligand-binding sites in other class C GPCRs, such as metabotropic glutamate receptors (mGluRs) and calcium sensing receptor (CaSR) ^33,34^. In addition, the binding of chloride ion in Tas1r3 and sodium ion in Tas1r1 was observed at the corresponding sites in medaka Tas1r2a/Tas1r3 ^11,35^ (Fig. 1A, 1B, 1D, Extended Data Fig. 3A). Although the ions were not modeled in the reported human TAS1R2/TAS1R3 structure, additional density blobs were observed at the corresponding sites for chloride and sodium ions in the density map of human TAS1R2/TAS1R3 ^24^ (Extended Data Fig. 3C, 3D), implying the presence of ions. These observations suggest that these ion-binding sites are conserved among TAS1Rs.

**Figure 1.**
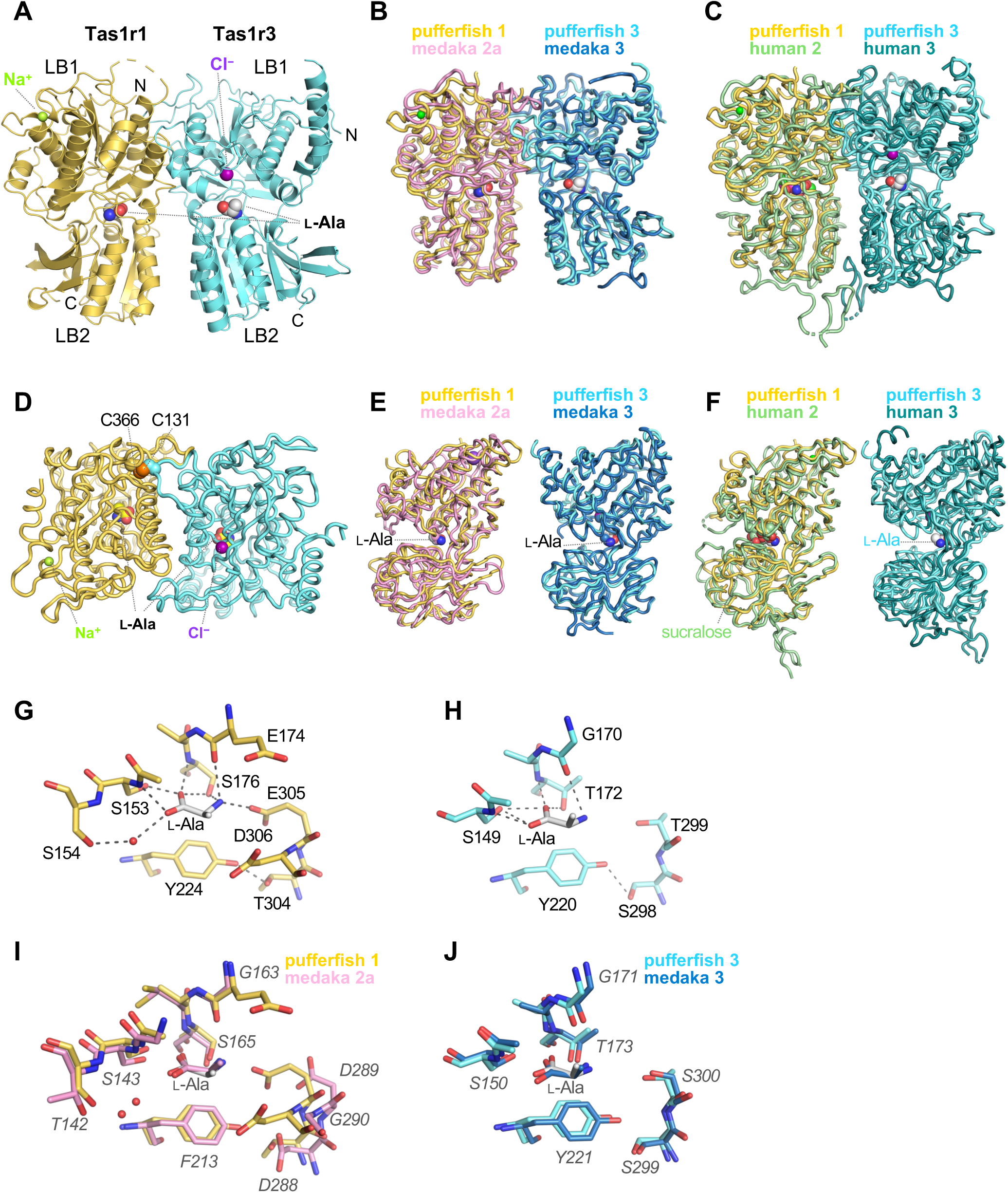
Structure of pufferfish Tas1r1/Tas1r3LBD. (A) Overall structure of pufferfish Tas1r1 (orange)/Tas1r3 (cyan)-LBD heterodimer. (B, C) Superposition of pufferfish Tas1r/Tas1r3LBD and medaka Tas1r2a/Tas1r3LBD (B; PDB ID: 5X2N) or human TAS1R2/TAS1R3 (C; PDB ID: 9NOV). (D) Inter-subunit disulfide bond in pufferfish Tas1r1/Tas1r3LBD viewed from the top of the panel A. (E, F) Subunit conformations in pufferfish Tas1r/Tas1r3LBD compared with those in medaka Tas1r2a/Tas1r3LBD (E; PDB ID: 5X2N) or human TAS1R2/TAS1R3 (F; PDB ID: 9NOV). (G, H) Close up views of the amino-acid binding site in Tas1r1 (G) and Tas1r3 (H). (I, J) Superposition of the amino-acid binding sites in pufferfish Tas1r1/Tas1r3LBD and medaka Tas1r2a/Tas1r3LBD. Amino acid residues in medaka Tas1r2a/Tas1r3The labels are labeled and italicized. (I) Pufferfish Tas1r1 (orange) and medaka Tas1r2a (pink). (J) Pufferfish (cyan) and medaka (blue) Tas1r3. In panels G–J, water molecules are shown as red dots.

In the substrate-binding site in pufferfish Tas1r1, the ligand L-alanine forms intermolecular interactions with the receptor, similar to those observed in medaka Tas1r2a and Tas1r3 (Fig. 1G, H). Specifically, the α-carboxyl group of L-alanine forms hydrogen bonds with the side-chain hydroxyl groups of Ser153 and Ser176; the α-amino group forms hydrogen bonds with the main-chain carbonyl group of Glu174; and the α-hydrogen interacts with Tyr224. Additionally, a salt bridge is formed between the α-amino group of the ligand L-alanine and the carboxyl group of Glu305. Amino acid binding was also observed at the corresponding site in Tas1r3, as observed in medaka Tas1r2a/Tas1r3, although Tas1r3 has been considered an auxiliary subunit in heterodimeric Tas1rs in terms of substrate recognition ^11^ (Fig. 1I, J). Here, the α-carboxyl group of L-alanine forms hydrogen bonds with the side-chain hydroxyl groups of Ser149 and Thr172, the α-amino group forms hydrogen bonds with the main-chain carbonyl group of Gly170, and the α-hydrogen makes a CH-π interaction with Tyr220. These structural configurations for amino acid recognition are conserved across pufferfish and medaka Tas1rs, as well as other class C GPCRs, such as mGluRs and CaSR.

The structural features observed in pufferfish Tas1r1/Tas1r3LBD are similar to those observed in medaka Tas1r2a/Tas1r3 and human TAS1R2/TAS1R3, and thus are likely conserved across TAS1Rs. Furthermore, the residues essential for ligand amino-acid recognition, which we previously termed as the “SSΩ” residues (Ser153, Ser176 and Tyr224 in pufferfish Tas1r1), are conserved among TAS1R1/TAS1R3s in various species, including the human umami receptor ^36^ (Extended Data Fig. 2). Therefore, the amino acid recognition mechanism observed in pufferfish Tas1r1/Tas1r3 is likely conserved across other TAS1R1/TAS1R3s.

### Pufferfish Tas1r1/Tas1r3 is a broad amino acid receptor with non-rigorous stereoselective recognition

To further investigate the structure-function relationship of pufferfish Tas1r1/Tas1r3, we examined the substrate specificity of this receptor by a ligand binding assay using Tas1r1/Tas1r3LBD in terms of protein thermal stabilization ^23^, as well as a receptor response assay using full-length Tas1r1/Tas1r3 in terms of cytosolic Ca^2+^elevation ^37^ (Fig. 2A, Extended Data Fig. 5A). The receptor bound and responded to a wide range of L-amino acids, including L-alanine. The broad L-amino acid specificity of pufferfish Tas1r1/Tas1r3 was similar to that of medaka Tas1r2a/Tas1r3 ^23^ and mouse TAS1R1/TAS1R3 ^2,9^, although different from that of human TAS1R1/TAS1R3, which is specific to glutamate or aspartate ^3^.

**Figure 2.**
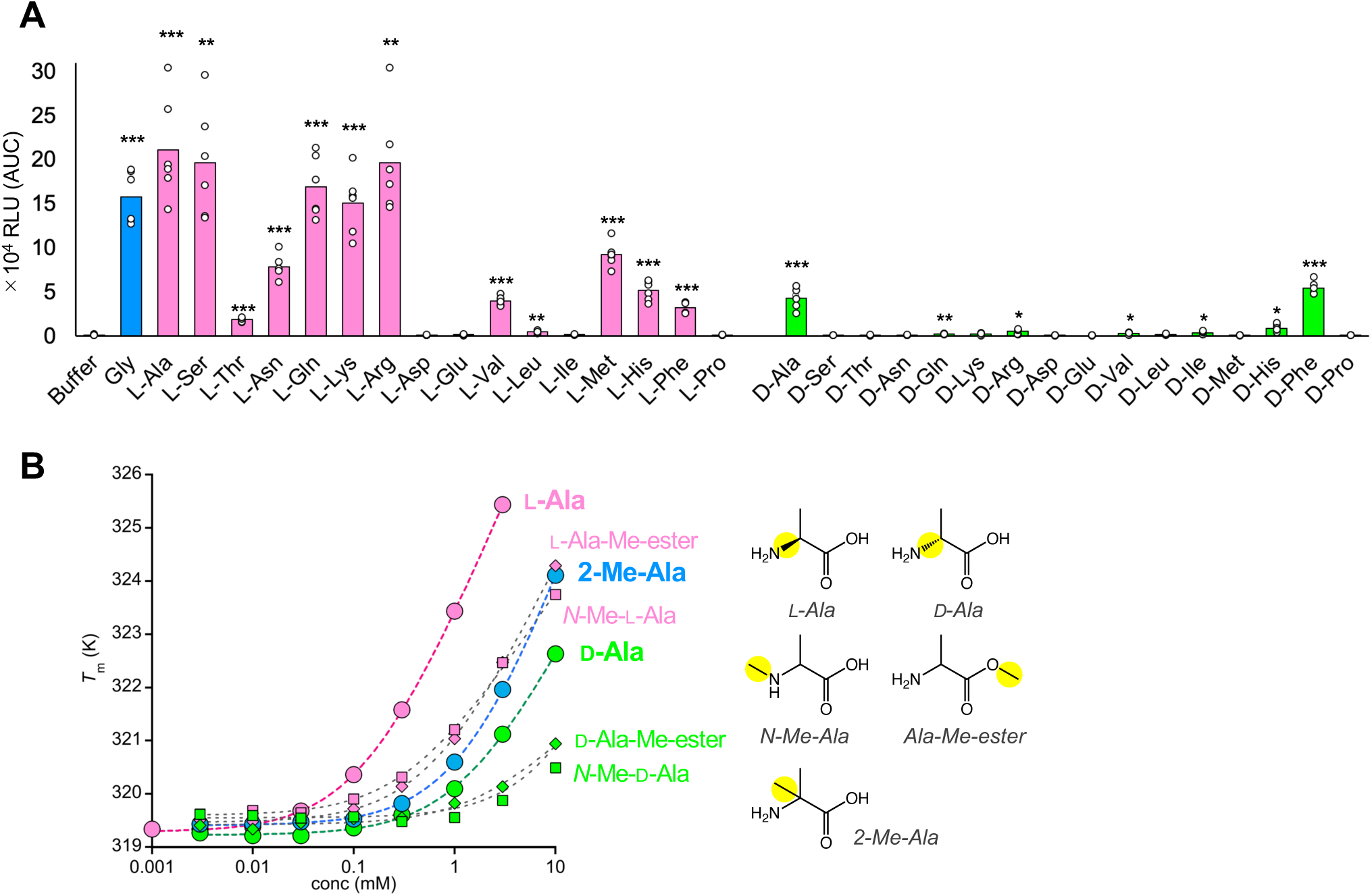
Substrate specificity of pufferfish Tas1r1/Tas1r3. (A) Responses of the full-length pufferfish Tas1r1/Tas1r3 evoked by 50 mM amino acids. Values are mean from six experiments. *, **, ***: Welch’s *t* test with Benjamini-Hochberg correction; \**p* < 0.05, \*\**p* < 0.01, \*\*\**p* < 0.001. (B) Dose-dependent thermal stabilization of Tas1r1/Tas1r3LBD induced by amino acid derivatives, analyzed by DSF.

Furthermore, pufferfish Tas1r1/Tas1r3LBD also exhibited binding to various D-amino acids (Extended Data Fig. 5B). Among these, D-alanine and D-phenylalanine demonstrated a relatively high extent of binding and evoked low but significant receptor responses (Fig. 2A). These observations are consistent with a previous report that showed taste nerve responses to D-alanine in pufferfish, albeit lower than those to L-alanine ^38^. The L-amino acid responsiveness of TAS1Rs is plausible because the content of free L-amino acids is significantly higher than that of D-enantiomers in most foods for animals according to protein homochirality. Nevertheless, the ability of D-amino acid sensing is also likely meaningful from the perspective of taste perception; for instance, substantial amounts of free D-amino acids are present in mollusks and crustaceans, the primary food of pufferfish (i.e., D-alanine constitutes 30–84% of total free alanine in crustaceans and Heterodonta mollusks as an osmolyte) ^39,40^. Canonically, L- and D-amino acids are thought to be recognized by a different set of TAS1R members; in humans and mice, L-amino acids are recognized by the umami receptors TAS1R1/TAS1R3s, while D-amino acids are recognized by the sweet receptors TAS1R2/TAS1R3s ^2,3^. Indeed, we confirmed that human TAS1R1/TAS1R3 showed high selectivity for L-glutamate over its D-enantiomer, eliciting only background level signals (Extended Data Fig. 3E). In the case of fish TAS1Rs, including medaka Tas1r2a/Tas1r3, binding and/or responses have been reported for L-amino acids, but not for D-amino acids, although a limited number of compounds have been tested for the latter ^22,23,28,36^. In contrast, pufferfish Tas1r1/Tas1r3 showed promiscuous stereochemical substrate recognition for both L- and D-amino acids.

To further address the substrate specificity of pufferfish Tas1r1/Tas1r3LBD, we examined its binding to amino acid derivatives. Chemicals lacking either the α-amino or α-carboxy groups of alanine (propionate or ethylamine) showed no obvious binding to the receptor, indicating their specificity for α-amino acids (Extended Data Fig. 5C). Although modifications to these functional groups (*N*-methylation or methyl esterification) showed moderately reduced binding, they did not abolish it (Fig. 2C, Extended Data Fig. 5C, Extended Data Table 2), as also observed for medaka Tas1r2a/Tas1r3 ^36^. However, unlike medaka Tas1r2a/Tas1r3, pufferfish Tas1r1/Tas1r3LBD also bound 2-methylalanine, an achiral alanine derivative with an additional methyl group at Cα instead of α-hydrogen, with an extent between those observed for L-alanine and D-alanine. Indeed, the apparent dissociation constants for L-alanine, D-alanine, and 2-methylalanine were estimated as 0.126 ± 0.007, 1.44 ± 0.09, and 1.29 ± 0.14 mM, respectively (Fig. 2C, Extended Data Table 2). This binding property distinctively contrasts with that of medaka Tas1r2a/Tas1r3, which showed substantially reduced binding of 2-methylalanine compared to L-alanine, that is, >100-fold increase in *K*_d_, indicating the significance of the CH-π interaction between the α-hydrogen in alanine and the aromatic residues immediately beneath the substrate for recognition ^36^. These results suggest that the insignificance of the CH-π interaction and/or a greater tolerance to the steric hindrance between the α-methyl group in 2-methylalanine and the aromatic residue underlies the non-rigorous enantiospecific amino acid recognition by pufferfish Tas1r1/Tas1r3.

Other than amino acids, no significant binding was observed for the other tested chemicals, including taste substances for humans, except for thermal destabilization caused by denatonium, which is unlikely to be related to receptor activation (Extended Data Fig. 5D). In addition, nucleotide- or betaine-induced enhancement of amino acid interactions, which have been reported in mammalian TAS1R1/TAS1R3 ^2,3^ and in pufferfish taste nerve responses ^31^, was not observed (Extended Data Fig. 5E).

### D-Amino acid-bound structures of pufferfish Tas1r1/Tas1r3LBD

To address the structural basis of non-rigorous enantioselective recognition, we determined the crystallographic structures of pufferfish Tas1r1/Tas1r3LBD in complex with D-amino acids that induced explicit receptor responses, namely D-alanine and D-phenylalanine (Extended Data Table 1). Difference Fourier analysis showed clear electron densities at the positions where L-alanine binds in each subunit in the cleft between LB1 and LB2 (Extended Data Fig. 4C–F). These results indicate that pufferfish Tas1r1/Tas1r3 recognize both L- and D-amino acids at the same sites.

The D-alanine-bound structure adopts a similar conformation as the L-alanine-bound structure (Fig. 3A), i.e., the compact dimerization state with the inter-subdomain cleft in both subunits “closed.” At the binding site, the D-alanine model is most favorably placed in a manner with keeping the α-amino and carboxy groups in the same position as L-alanine, while flipping the α-methyl group and α-hydrogen, as determined by electron density maps (Fig. 3B, 3C, Extended Data Fig. 4C, D). Indeed, NMR analysis of ^13^C^15^N-alanine binding to the pufferfish Tas1r1/Tas1r3LBD protein showed that the α-methyl groups in L- and D-alanine are under different chemical environments at the binding sites (Extended Data Fig. 4G, H): the ^13^C-edited NOESY peaks relevant to the α-methyl groups resulted in different correlation patterns between L- and D-enantiomers. Thus, the hydrogen-bonding network observed in the recognition of L-alanine was also conserved in D-alanine recognition. A remarkable change was observed on the interaction with the aromatic residues immediately beneath the substrate, Tyr224 in Tas1r1 and Tyr220 in Tas1r3: the CH-π interactions with L-alanine are now replaced with the hydrophobic interaction with the methyl group in D-alanine. This interaction switching agrees with the NMR spectra showing differences in the NOE signals, presumably between the methyl proton in L- or D-alanine and the aromatic ring (Extended Data Fig. 4G, H). This observation is also consistent with the binding assay results showing that 2-methylalanine exhibited a weaker interaction than L-alanine but remained able to bind to pufferfish Tas1r1/Tas1r3LBD (Fig. 2B, Extended Data Fig. 5C), but not significantly to medaka Tas1r2a/Tas1r3LBD ^36^. These results again indicated the non-rigorousness of the CH-π interaction between the α-hydrogen in the amino acid and the aromatic residue in the receptor, despite the fact that all essential residues for recognition of amino acids in L-enantiospecific TAS1Rs, including the aromatic residue, are conserved in this receptor.

**Figure 3.**
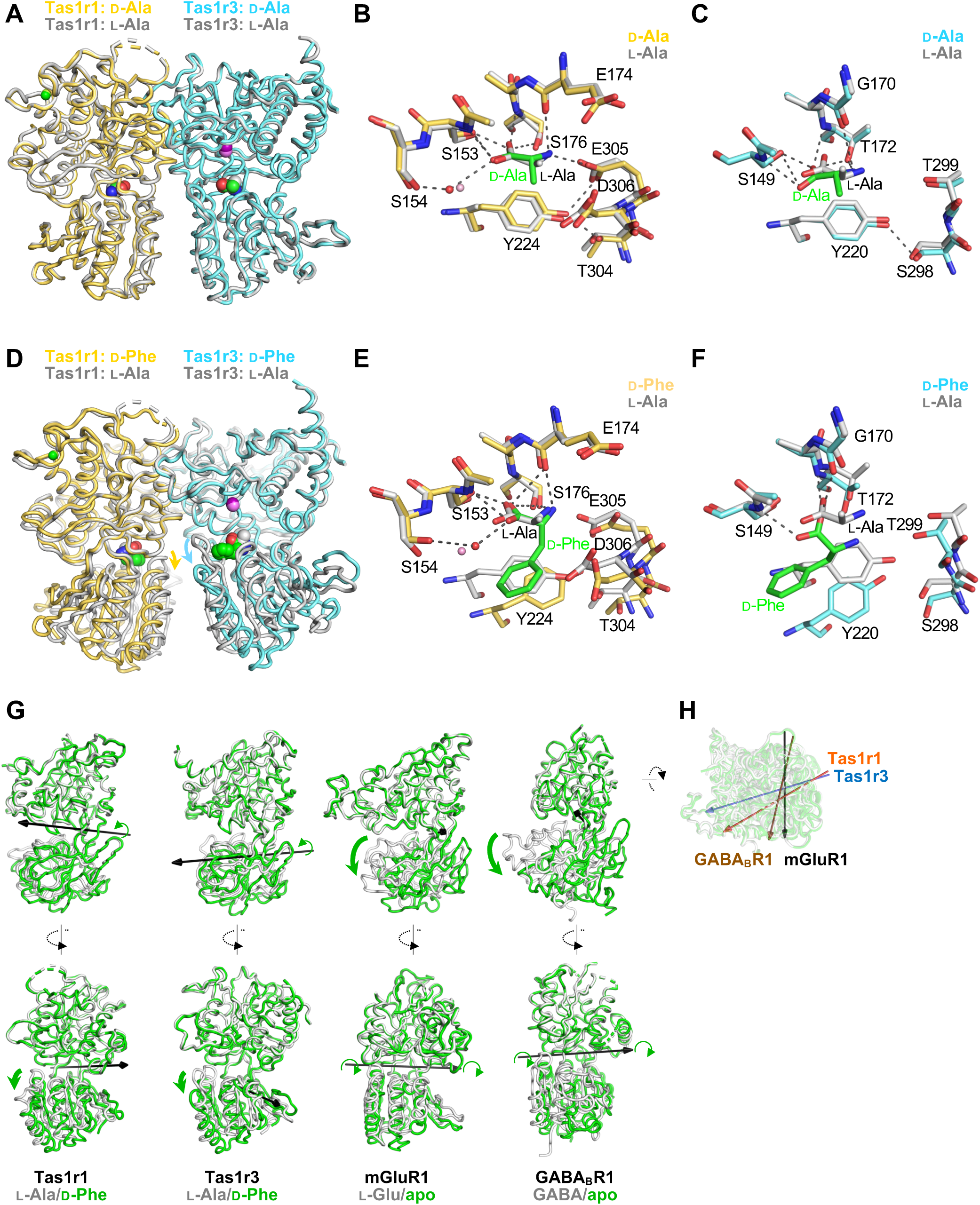
D-amino acid recognition of pufferfish Tas1r1/Tas1r3. (A–C) D-Alanine-bound Tas1r/Tas1r3LBD structure. D- (yellow and cyan) and L- (gray) alanine-bound Tas1r/Tas1r3LBD structures were superposed. (A) Overall structures. (B, C) Close up views of the amino-acid binding site in Tas1r1 (B) and Tas1r3 (C). (D–F) D-Phenylalanine-bound Tas1r/Tas1r3LBD structure. D-phenylalanine (yellow and cyan) and L-alanine (yellow)- bound Tas1r/Tas1r3LBD structures were superposed. (D) Overall structures. (E, F) Close up views of the amino-acid binding site in Tas1r1 (E) and Tas1r3 (F). (G, H) Domain motion analysis of LBDs of class C GPCRs. Side, front (G) and top (H) views of LBDs with arrows representing the rotation axes of domain motions between two states with the different ligands shown at the bottom of each panel. The domain motions were analyzed by DynDom ^63^.

The D-phenylalanine-bound Tas1r1/Tas1r3LBD structure also adopts a compact dimerization manner, similar to that observed for the other structures, but with a noteworthy difference in each subunit conformation (Fig. 3D). Within the binding site, D-phenylalanine is recognized analogously to D-alanine, orienting its α-substituent group facing the aromatic residues Tyr224 in Tas1r1 and Tyr220 in Tas1r3 (Fig. 3E, 3F). However, the phenyl group of D-phenylalanine forms a π-π interaction with the hydroxyphenyl groups of these tyrosine residues in the receptor. This interaction pushes the side chain downward, causing a subsequent downward shift in LB2, in which these tyrosine residues exist. Consequently, this resulted in a slight gap opening in the cavities (Fig. 3D). Nevertheless, the openings in the D-phenylalanine-bound Tas1r1 and Tas1r3 structures were substantially narrower than those in the so-called “open” conformations observed in the apo- or antagonist-bound LBD structures of other class C GPCRs, such as mGluRs, GABA_B_R, and CaSR (Fig. 3G, 3H, Extended Data Fig. 6A, Extended Data Table 3). Furthermore, the opening mechanism differs from the typical hinge-like motion observed in other class C GPCRs: the rotational axes run laterally across the hinge between LB1 and LB2. In contrast, the motion in pufferfish Tas1r1 and Tas1r3 involves rotational axes running longitudinally along the right-hand side of the gap opening. As a result, the structural configuration on this pivotal side of each subunit is mostly retained in its “closed” state, even with the gap present.

### Inter-subdomain interactions in TAS1R-LBDs and enantioselectivity

Which structural factors in the pufferfish Tas1r1/Tas1r3LBD enable a “closed” conformation despite the presence of a gap? In Tas1r1, we identified two “latch-like” inter-subdomain interactions contributing to cleft closure: hydrogen bonding between Tyr385 (LB1) and Asp306(LB2) proximal to the bound amino acid (Fig. 4A) and a cation-π interaction between Tyr332 (LB1) and Arg469 (LB2) distal to the bound amino acid (Fig. 4B). These interactions are located on the right side of the bound amino acids. In particular, the distal latch retained the interaction even in D-phenylalanine-bound Tas1r1, which is responsible for closing the cleft with a gap (Fig. 3D, 3G). In Tas1r3, a salt bridge and a hydrogen bond were formed between Arg385 (LB1) –Glu469 (LB2) and Gln325 (LB1) – Asn474 (LB2), respectively, in the distal regions of the bound amino acid (Fig. 4C).

**Figure 4.**
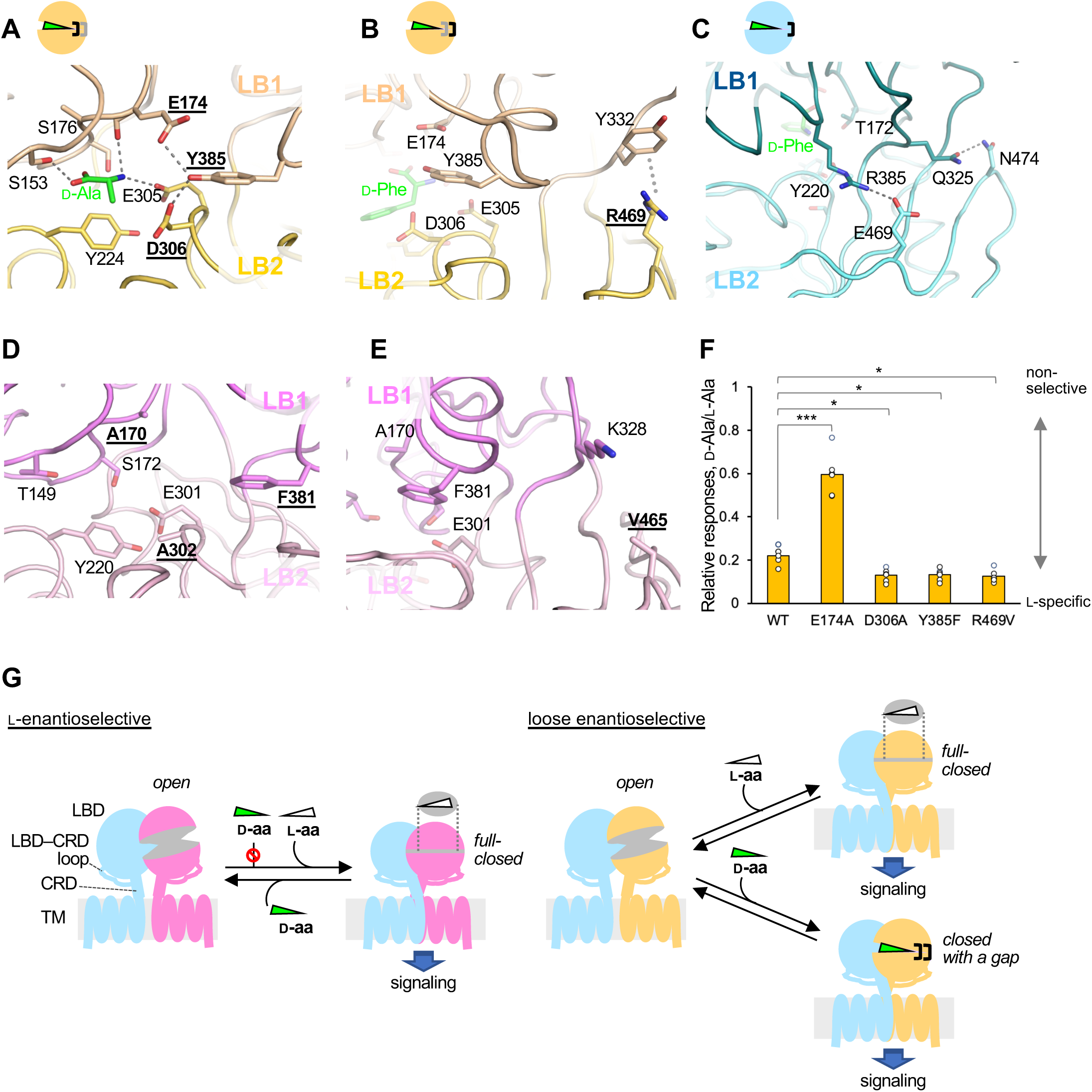
Inter-subdomain interactions and enantioselectivity of pufferfish Tas1r1/Tas1r3. (A, B) Inter-subdomain interactions in pufferfish Tas1r1 at the proximal (A) and the distal (B) sites to the bound amino acids. (C) Inter-subdomain interactions in pufferfish Tas1r3. (D, E) The corresponding sites in the human TAS1R1 structure predicted by AlphaFold3 ^65^ to those shown in panel A and panel B. The residues corresponding to those forming the “latch-like” interactions in pufferfish Tas1r1 are in bold and underlined. (F) d-Alanine responses relative to l-alanine responses of pufferfish Tas1r1/Tas1r3 and their mutants with a disruption/enhancement of the “latch-like” interactions. Displayed are relative d-alanine/l-alanine responses at 50 mM by Tas1r1-mutant/Tas1r3-wild-type receptors (Supplementary Fig. 1) with mutations introduced in Tas1r1 shown at the bottom. Values are mean from six experiments. *, **, ***: a two-sided one-way analysis of variance with Dunnett’s test; \**p* < 0.05, \*\**p* < 0.01, \*\*\**p* < 0.001. (G) Speculative schematic drawing of receptor activation of l-enantioselective and loose enantioselective TAS1Rs.

We then examined the relevance of these inter-subdomain “latch-like” interactions observed in the pufferfish Tas1r1/Tas1r3 structure to the loose enantioselectivity of this receptor. Notably, the specific inter-subdomain interactions found in pufferfish Tas1r1 were not conserved in the crystal structure of medaka Tas1r2a (Extended Data Fig. 6B, C) or in the predicted structure of human TAS1R1 (Fig. 4D, E). While medaka Tas1r3 exhibits similar, though not identical, inter-subdomain interactions in its “closed” structure (Extended Data Fig. 6D), non-TAS1R3 subunits are generally considered the primary determinant of substrate specificity ^41^. Therefore, we focused on the Tas1r1 subunit in the pufferfish Tas1r1/Tas1r3. We mutated Tyr385, Asp306, and Arg469 (i.e., residues forming the “latch-like” interactions) with the corresponding amino acids in human TAS1R1, a representative L-specific ortholog (Fig. 4E). Additionally, we tested the mutation at Glu174, which may have formed a hydrogen bond with Tyr385, competing with the interaction between Tyr385 and Asp306, because both Glu174 and Asp306 serve solely as hydrogen-bond acceptors (Fig. 4A).

A ligand-binding assay using the purified Tas1r1/Tas1r3LBD variants confirmed that the amino acid affinities of these mutants were largely unchanged; the *K*_d_ values were within two-fold of those of the wild-type (Extended Data Table 4). This is consistent with the structural observation that the side chains of these residues do not directly interact with bound amino acids. However, analysis of full-length mutant signaling revealed significant differences in the responses to D-alanine compared to the wild-type receptor (Fig. 4F and Supplementary Fig. 1). We evaluated the relative responses induced by 50 mM D-alanine compared with those induced by 50 mM L-alanine, which induced near-maximal responses in all receptors (Supplementary Fig. 1). Mutants disrupting LB1-LB2 interactions (D306A, Y385F, and R469V) showed a reduced response to D-alanine and thus became more L-enantioselective. Although the reduced response observed in the D306A mutant could partly be attributed to the reduction in D-alanine binding affinity, the D-alanine-specific reduction in the Y385F and R469V mutants occurred despite unchanged binding affinities (Extended Data Table 4). This decoupling of the binding affinity and receptor response indicates that these “latch-like” interactions do not contribute to the initial ligand binding but rather that they contribute to the efficacy of the subsequent conformational transition toward the active state.

In sharp contrast, the E174A mutant exhibited a significantly increased response to D-alanine, resulting in a loss of L-enantioselectivity, despite the unchanged D-alanine binding affinity. The E174A mutation disrupts a hydrogen bond with Tyr385, which is an alternative to, and thus competes with, the formation of the LB1-LB2 inter-subdomain hydrogen bond between Tyr385 and Asp306. Therefore, E174A likely enhances intersubdomain interactions. In summary, these observations indicate that the modulation of LB1-LB2 interactions in Tas1r1 dictates the enantioselective responses of the receptor.

These results suggest a plausible mechanism underlying non-rigorous stereospecific recognition of this receptor (Fig. 4G). Pufferfish Tas1r1 possesses the “latch-like” inter-subdomain interactions on the right side of the amino acid-binding cleft. These interactions stabilize the closed conformation, maintaining the local protein structure, even when a ligand fits imperfectly into the binding site, resulting in a gap. Because the C-terminus of the LBD, which connects to the downstream structural domain as the cysteine-rich domain (CRD), is located on the right side of the LBD toward the cleft (Fig. 1A), this stabilization by the latches likely facilitates conformational changes in the further downstream transmembrane region required for receptor activation and signal transduction, even with suboptimal ligand binding (Fig.4G, bottom right). In contrast, L-enantioselective receptors lacking such inter-subdomain interactions may have been unable to adopt the fully closed LBD conformation required for signaling, even if a D-amino acid bound (Fig. 4G, left).

## Discussion

Taste has the functional requirement of recognizing a wide variety of chemicals in food with a limited repertoire of receptors. In the case of TAS1Rs, which are responsible for the sensation of organic nutrients and thus for promoting animal eating, the receptors are likely diversified in two respects, relating to the diversification of the food habits of animals: diversification into multiple different subtypes and acquisition of a different chemical specificity in each subtype. The crystallographic structure of the pufferfish Tas1r1/Tas1r3LBD provides insights into both aspects. First, the study verified that the basic protein architectures of TAS1Rs, at least in LBDs, are conserved across different subtypes. Second, the study indicated that a different chemical specificity can be achieved not by a gain or loss of direct interactions with the ligands, but rather by a gain or loss of intramolecular inter-subdomain interactions within the receptor, conferring a different conformational equilibrium upon ligand binding (Fig. 4G).

A previous study has suggested that the broad amino acid specificity of mouse TAS1R1/TAS1R3 is attributed to the residues with no direct interaction with the ligand amino acid in comparison with human TAS1R1/TAS1R3, which is specific to L-glutamate or aspartate: most of the mutations conferring a broad responsiveness to human TAS1R1/TAS1R3 are located at the periphery of the cleft and the interface between LB1 and LB2 in TAS1R1 ^9^. These mutations are considered to alleviate charge repulsion with the surroundings and thus enhance inter-subdomain interactions, similar to the E174A mutant in pufferfish Tas1r, which enhances D-amino acid responses. Therefore, a broad substrate specificity in mouse TAS1R1 and pufferfish Tas1r1 with a further broadness across the stereochemistry was likely achieved by similar structural contexts, despite the fact that the responsible regions are not identical. In addition, since E174 is in the vicinity of the bound amino acid, a charge modification by the alanine mutation could also affect substrate specificity, similar to the E171A mutation in mouse TAS1R acquired L-glutamate binding ability ^9^. In addition to the “latch-like” inter-subdomain interactions, the D-phenylalanine-bound pufferfish Tas1r1/Tas1r3LBD exhibited the specific π– π interactions between the phenyl groups in the ligand and the side chains of the aromatic residues immediately beneath the ligand, also retaining cleft closure. Intriguingly, human and murine sweet taste receptors TAS1R2/TAS1R3 show higher responses to aromatic D-amino acids such as D-phenylalanine, among other D-amino acids ^2,3,42^. Of note, no “latch-like” interactions at the sites corresponding to those found in pufferfish Tas1r1/Tas1r3 were observed in human TAS1R2/TAS1R3 (Extended Data Fig. 6E-G). Since the D-amino acid-bound structure of the human sweet receptor has not yet been reported, future studies should address whether the mechanisms of D-amino acid recognition by the sweet receptors are similar to those found in pufferfish Tas1r1/Tas1r3.

Macromolecules are, in many cases, stereochemically rigorous and recognize and/or produce only one enantiomer. In the case of enzymes, usually only one enantiomer of a reactant, which is preferentially placed at the active site, serves as the substrate ^43^. Stereochemical rigorousness for substrate recognition has also been observed for receptor proteins, such as amino acid receptors, including ionotropic and metabotropic GluRs and CaSR ^44-47^, with some rare exceptions, such as the NMDA receptor, which can accommodate both L- and D-aspartate and their derivatives without a steric clash ^48^. In fact, taste sensation is the first example in which stereoselectivity at the receptor level in humans was discovered 140 years ago ^49,50^. In the case of amino acid sensation, while the major free amino acids in organisms serving as food are proteinogenic L-amino acids, it may also be meaningful to sense D-amino acids, as some aquatic invertebrates are rich in D-alanine ^39,40^. Indeed, marine invertebrate-eating fish are known to have high D-amino acid oxidase activity in their intestines ^50^, suggesting the existence of a metabolic system for D-amino acids. However, it remains unclear how stereospecifically rigorous the other TAS1Rs are, or whether some TAS1Rs are as stereochemically promiscuous as pufferfish Tas1r1/Tas1r3. For example, mouse TAS1R1/TAS1R3 is specific to L-amino acids but responds to D-alanine in the presence of nucleotides ^2^, while human TAS1R2/TAS1R3 is specific to D-amino acids, but several L-amino acids are known to evoke sweet sensations ^51^, although the responses have not been proven at the receptor level ^42^. Therefore, extensive analyses of the substrate specificities of TAS1Rs, including their stereospecificities, are required.

Previous studies have identified several strategies for shaping the substrate specificity of TAS1Rs. For example, L-glutamate specificity found in TAS1R1/TAS1R3 in humans and other leaf-eating primates can be explained by the replacement of acidic residues with neutral amino acids to avoid charge repulsion ^6,9^, and the inosine monophosphate sensitivity in TAS1R1/TAS1R3s is obtained by the introduction of basic residues providing ionic interactions with the molecule close to its binding site ^52^. This study structurally elucidates a distinct strategy to extend the range of detectable chemicals, not by introducing a direct interaction with the chemical, but rather by introducing intramolecular interactions to modulate the conformational equilibrium of the receptor when the chemical binds.

## Methods

### Protein preparation

To construct the expression vector for *T. rubripes* Tas1r1LBD, pAc_fft1r1L_P, the gene encoding the LBD region of Tas1r1 (Met1–Ser501), followed by the PA-tag (GVAMPGAEDDVV), was subcloned into pAc5.1/V5-HisA (Invitrogen) between the *Kpn*I and *Pme*I sites. To construct the expression vector for *T. rubripes* Tas1r3LBD, pAc_fft1r3L_F, the gene encoding the LBD region of Tas1r3 (Met1–Ser496), followed by the FLAG-tag (DYKDDDDK), was subcloned in the same manner as pAc_fft1r1L_P. Subsequently, to eliminate free cysteines at the surface, Cys242 in Tas1r1 and Cys238 in Tas1r3, which are expected to form disulfide bonds with downstream cysteine residues in the full-length proteins but remain free in the isolated LBDs, were mutated to serine by polymerase chain reaction (PCR) using PrimeSTARMax (Takara Bio), resulting in the vectors pAc_fft1r1L_C242S_P and pAc_fft1r3L_C238S_F. These vectors are treated as the “wild-type” for convenience, because the introduced surface mutations are distant from the active sites (Extended Data Fig. 1G) and there are no mutations in the vicinity of the active sites. For Tas1r1, the E174A, D306A, Y385F, and R469V mutations were introduced into the pAc_fft1r1L_C242S_P vector by PCR. The expression vectors of wild-type or mutant Tas1r1 and wild-type Tas1r3, together with the antibiotic marker vector pCoBlast (Invitrogen), were co-introduced into *Drosophila* S2 cells (Invitrogen) using the calcium phosphate transfection method or the lipofectant reagent X-tremeGene HP (Roche), and a high-expression stable cell clone was established by selection using the expression levels of both Tas1r1LBD and Tas1r3LBD genes as indicators, as described previously ^28^. The genomic sequence of each cell clone was analyzed to verify the introduction of Tas1r coding sequences and mutations at the corresponding sites in Tas1r1.

The established S2 cells, which stably express the C-terminal PA-tagged Tas1r1LBD (C242S mutant) and the C-terminal FLAG-tagged Tas1r3LBD (C238S mutant), were cultured in 90% Schneider’s Drosophila Medium (Gibco), 10% Fetal Bovine Serum, 0.025% Pluronic F-68 (Gibco), or that mixed with ExpressFiveSFM (Gibco) at a 1:1 ratio at 27 °C in the presence of 10 µg/mL Blasticidin S. Seven days prior to protein purification, the cells were centrifuged and resuspended in the fresh medium with the same composition as described above, with the exception of Blasticidin S, and cultured further at 20 °C.

Tas1r1/Tas1r3LBD was purified from the culture medium using ANTI-FLAG M2 Affinity Gel (SIGMA-Aldrich) at 4 °C, as described previously ^21^. Briefly, after the culture medium was collected by centrifugation, phenylmethylsulfonyl fluoride, L-alanine, 2 M Tris-HCl solution (pH 8.0), and CaCl_2_ were added at final concentrations of 0.5 mM, 0.1 M, 40, and 40 mM, respectively. After incubation for 60 min, the medium was centrifuged, filtered, and applied to the ANTI-FLAG M2 Affinity Gel. The FLAG resin was washed with 25 to 40 column volumes of buffer A (20 mM Tris, 0.1 M L-alanine, 2 mM CaCl_2_, 0.3 M NaCl, pH 8.0), and the protein was eluted with 5 column volumes of buffer A containing 100 µg/mL FLAG peptide. The protein concentration was estimated by absorbance at 280 nm with a Nanodrop (Thermo Scientific) using the extinction coefficient calculated from the amino acid sequences (Abs_280nm_[1 mg/mL] = 1.538). The protein samples were stored at 4 °C until use.

### Structural analysis

The “wild-type” Tas1r1/Tas1r3LBD protein (free-cysteine eliminated; Tas1r1- C242S/Tas1r3- C238S) sample was prepared as described in the section “Protein preparation,” except that Kifunensine was added to 5 µM to the culture medium at the stage of medium exchange seven days before purification. The purified sample was mixed with Endo H at a ratio of 0.05 mg of Endo H to 1 mg of T1r1/T1r3LBD, and incubated at 4 °C overnight to remove the glycosyl chains. The sample was diluted with buffer B (20 mM Tris-HCl, 0.1 M L-alanine, 2 mM CaCl_2_, pH 8.0) to reduce the NaCl concentration below 50 mM and subjected to anion exchange chromatography using Resource Q 1mL (GE Healthcare) connected with an Äkta purifier (GE Healthcare). The protein was eluted using an NaCl gradient from 50–300 mM through 40 column volumes. The protein was eluted mainly in two peaks: peak A at a conductivity of 19.8 mS/cm and peak B at 22.5 mS/cm (Extended Data Fig. 1A–C). The fractions at peaks A and B were collected separately and confirmed to consist of the pufferfish T1r1/T1r3LBD heterodimer by N-terminal sequencing. The sample was concentrated and buffer-exchanged to buffer C (10 mM Tris-HCl, 0.1 M L-alanine, 2 mM CaCl_2_, 50 mM NaCl, pH 8.0) using Vivaspin-6 (MWCO: 10,000, Sartorius).

L-Alanine-bound Tas1r1/Tas1r3LBD crystals were obtained using the sitting-drop vapor diffusion method. The protein solution from the peak B fraction (∼4 mg/mL) was mixed with 0.1 M MES-NaOH (pH 6.0), 18–22% PEG3350, and 20% glycerol and incubated at 20 °C. D-Alanine-bound T1r1/T1r3 was prepared in the same manner as the L-alanine-bound sample, except that the L-alanine in buffers A, B, and C was replaced with D-alanine. The D-alanine-bound Tas1r1/Tas1r3LBD crystals were obtained under the same conditions as used to obtain the L-alanine-bound crystals. For D-phenylalanine-bound Tas1r1/Tas1r3 preparation, the protein sample was prepared in the same manner as the L-alanine-bound sample, except that 0.1 M L-alanine in buffers B and C was replaced with 30 mM D-phenylalanine. The D-phenylalanine-bound Tas1r1/Tas1r3LBD crystals were obtained under the same conditions as used to obtain the L-alanine-bound crystals, except that the protein sample from the peak A fraction was used for crystallization. The L- and D-alanine-bound peak A and D-phenylalanine-bound peak B fractions did not yield crystals with suitable qualities, even under similar crystallization conditions. The crystals were flash-frozen in liquid nitrogen before data collection.

The X-ray diffraction data were collected at BL32XU at SPring-8 at a wavelength of 1.0 Å using an EIGER X9M detector (DECTRIS). Because the obtained crystals were either small or highly clustered (Extended Data Fig. 1D–F), the diffraction data for each sample were collected from multiple crystals using the ZOO system ^53^ with the strategy of small-wedge synchrotron crystallography ^54^, and were merged with the KAMO system ^55^ based on intensity-based hierarchical clustering analysis ^56^.

The crystal structure of L-alanine-bound Tas1r1/Tas1r3LBD was solved by molecular replacement with MOLREP ^57^ using the predicted structures of pufferfish Tas1r1LBD and Tas1r3LBD constructed by AlphaFold2 ^58^ as a search model. The initial model was subjected to jelly body refinement using REFMAC5 ^59^. The model was manually rebuilt using COOT ^60^ and refined using Phenix ^61^. The crystal structures of D-alanine- and D-phenylalanine-bound Tas1r1/Tas1r3LBD were solved by molecular replacement with Phaser ^62^, using the L-alanine-bound Tas1r1LBD and Tas1r3LBD structures as search models. For the D-phenylalanine-bound structure, the initial model was further subjected to rounds of jelly body refinement. The models were further refined using Phenix, followed by manual rebuilding using COOT.

Conformational changes in Tas1r1/Tas1r3LBD and the LBDs of other class C GPCRs were analyzed using DynDom ^63^. Orientational differences in the rotation axes for inter-subdomain cleft opening/closure were evaluated as follows: two LBD structures of a receptor in different conformations were submitted to DynDom, and the output coordinates were superposed onto the L-alanine-bound Tas1r1LBD structure solved in this study. Then, the coordinates of the rotation axis derived from each DynDom analysis were converted to a vector, and the angle with the reference vector, which was derived from the rotation axis of the conformation change between the agonist-bound and apo mGlu1 structures ^34^, was determined.

### Ligand binding assay

A ligand binding assay using the “wild-type” (free-cysteine eliminated; Tas1r1- C242S/Tas1r3- C238S) and mutant pufferfish T1r1/Tas1r3LBD introduced on this “wild-type” backbone was performed by differential scanning fluorimetry (DSF), as described previously ^23^. Briefly, the purified protein was dialyzed against assay buffer (20 mM Tris-Cl or HEPES-NaOH, 300 mM NaCl, 2 mM CaCl_2_, pH 8.0). Protein samples (1 μg) were mixed with Protein Thermal Shift Dye (at 1× concentration; Applied Biosystems) and various concentrations of each ligand in 20 μL of the assay buffer, and loaded onto a MicroAmp Fast Optical 48-Well Reaction Plate (Applied Biosystems). The fluorescence Intensity was measured using a StepOne Real-Time PCR System (Applied Biosystems). The temperature was raised from 25 °C to 99 °C at a rate of 0.022 °C /s. The reporter and quencher for detection were set to ROX and none, respectively. The apparent melting temperature (*T*_m_) was determined using the peak of the derivative curve of the melting curve (dFluorescence/dT) using Protein Thermal Shift Software version 1.3 (Applied Biosystems). The apparent *K*_d_ (*K*_d-app_) was determined by fitting the results to an equation based on the simple thermodynamic folding model proposed by Schellman ^64^, as described previously ^23^.

### Luminometric receptor response assay

The responses of pufferfish and human TAS1R1/TAS1R3 were measured using heterologous expression systems, as previously described ^2,3,9^. HEK293T cells were transiently co-transfected with expression vectors for TAS1R1, TAS1R3, rat G15i2, and mt-apoclytin-II using Lipofectamine 2000 (Invitrogen). After 48 h, the transfected cells were trypsinized, seeded in 96-well black-walled CellBIND surface plates (Corning), and cultured overnight at 37 ℃ in 5% CO_2_. After overnight culture, the medium was removed and replaced with coelenterazine loading buffer (10 μM coelenterazine, 10 mM HEPES, 130 mM NaCl, 10 mM glucose, 5 mM KCl, 2 mM CaCl_2_, and 1.2 mM MgCl_2_, 0.1% BSA, pH adjusted to 7.4 using NaOH) for 4 h at 27 ℃ in the dark. After 20 s of baseline reading, an aliquot of the assay buffer supplemented with 2× ligand was added, and light emission was recorded using a FlexStation 3 microplate reader (Molecular Devices) for an additional 90 s. The response from each well was calculated based on the area under the curve (AUC) and expressed as relative light units (RLU). Statistically significant increases compared to the response induced by the buffer addition were determined using Welch’s two-sided *t*-test, and the Benjamini-Hochberg adjustment was used to correct for multiple comparisons (*q* = 0.05) using R (https://www.r-project.org/). In Fig. 4F, the RLU values after 50 mM D-alanine addition relative to those after 50 mM L-alanine addition are displayed, and the results of a two-sided one-way analysis of variance with Dunnett’s test with the wild-type as a control (KaleidaGraph, Synergy Software) are shown.

### NMR experiments

Purified Tas1r1/Tas1r3 LBDs were buffer-exchanged against an NMR buffer containing 20 mM Tris (hydroxymethyl-d_3_) amino-d_2_-methane-DCl (pH* 8.0), 300 mM NaCl, 2 mM CaCl_2_ and 1 mM [2,3-^13^C_2_; ^15^N]-D-alanine or [2,3-^13^C_2_; ^15^N]-L-alanine using PD-10 (Cytiva). The Water in the NMR buffer was 100% D_2_O. The concentration of the Tas1r1/Tas1r3 LBD samples ranged from 20–50 µM.

NMR measurements were performed using an Avance III HD 800 spectrometer equipped with a TXI cryogenic probe (Bruker Biospin) at 25 °C. The ^13^C-edited NOESY experiment was performed for intermolecular NOE analysis of the complex of each Tas1r1/Tas1r3 LBDs with [2,3-^13^C_2_; ^15^N]-D-alanine or [2,3-^13^C_2_; ^15^N]-L-alanine with a NOE mixing time of 200 ms. The data size and spectral width were 256 (*t1*) × 1(*t2*) × 2,048(*t3*) points and 11,160 Hz (*ω*_1_, ^1^H) × 10,060 Hz (*ω*_2_, ^13^C) × 11,160 Hz (*ω*_3_, ^1^H), respectively. The carrier frequencies of ^1^H and ^13^C were 4.7 and 35 ppm. The number of scans/FID was 256 (for 50 µM T1r1/T1r3LBD - 1 mM [2,3-^13^C_2_; ^15^N]-D-alanine complex), 512 (for 20 µM T1r1/T1r3LBD - 1 mM [2,3-^13^C_2_; ^15^N]-L-alanine complex), and 280 (40 µM T1r1/T1r3LBD - 1 mM [2,3-^13^C_2_; ^15^N]-L-alanine complex).

## Data Availability

Nucleotide sequences for *tas1r1* and *tas1r3* from *T. rubripes* used in this study have been deposited in the DNA Data Bank of Japan under the accession numbers LC843429 (*tas1r1*) and LC843430 (*tas1r3*). Coordinates and structure factors for the Tas1r1/Tas1r3LBD crystals have been deposited in the Protein Data Bank under the accession numbers 9XPJ (L-alanine-bound), 9XPK (D-alanine-bound), and 9XPL (D-phenylalanine-bound). Coordinates with the PDB IDs 5X2N, 9NOV, 9NOU, 1EWK, 1EWT, 1ISS, 6N51, 6N52, 7MTS, 7MTQ, 4MS3, 4MQE, 4MR7, 6UO8, 6VJM, 7C7Q, 7C7S, 5K5S, and 5K5T; and cryo-EM density map with the EMDB ID EMD-49167 were used in this study.

## Acknowledgement

We thank Professor Sadao Kiyohara for their valuable discussions on the taste responses of pufferfish; Drs. Yuko Kusakabe, Marina Kose, and Akihiro Itoigawa for conducting the preliminary response assays; and Yuri Kuhara, Keiko Hiratomi, Noriko Matsuura, and Fumie Iwabuki for technical assistance. The authors are grateful to Ms. Tsugumi Shiokawa and Dr. Hiroko Tada at Division of Instrumental Analysis, Okayama University, for the amino acid sequence analyses. Synchrotron radiation experiments were performed at SPring-8 with the approval of the Japan Synchrotron Radiation Research Institute (JASRI) (Proposal No. 2015A1795, 2018A1534, 2019A2539, 2020A2581, 2022A2719). This work was financially supported by JSPS KAKENHI Grant Numbers JP18H04621, JP20H04778, JP20H03195, JP23H02424, JP23K27117, JP24K21272 (to A.Y.), JP23K26861, and JP25H01362 (to Y.T.), the JST FOREST Program Grant Number JPMJFR220C (to Y.T.), Society for Research on Umami Taste (to A.Y.), Takeda Science Foundation (to A.Y.), OU Master Plan (to A.Y.), Platform Project for Supporting Drug Discovery and Life Science Research (Basis for Supporting Innovative Drug Discovery and Life Science Research [BINDS]) from AMED under Grant No. JP21am0101070 (support number 0585), JP23am121001 (support number 4929), and Lotte Shigemitsu Prize (to Y.T. and Y.I.). We used Paperpal by Editage (www.editage.jp) for English language editing and would like to thank their in-person human editing service.

## Author contributions

A.Y. conceived the study. R.M., M.N., T.Y., M.H., Y.A., A.Y., C.I., and N.Y. performed recombinant protein preparation. R.M., T.Y., M.H., and A.Y. performed crystallization. R.M., H.M., K.H., A.Y., and N.T. performed X-ray data collection and structure analysis. R.M., M.N., T.Y., and A.Y. performed the ligand binding assay. Y.T. and Y.I. performed the receptor response assay. Y.M. performed the NMR analysis. A.Y., Y.T., Y.M., T.Y., and R.M. wrote the paper, together with input from all of the other authors.

---

**Extended Data Figure 1.**
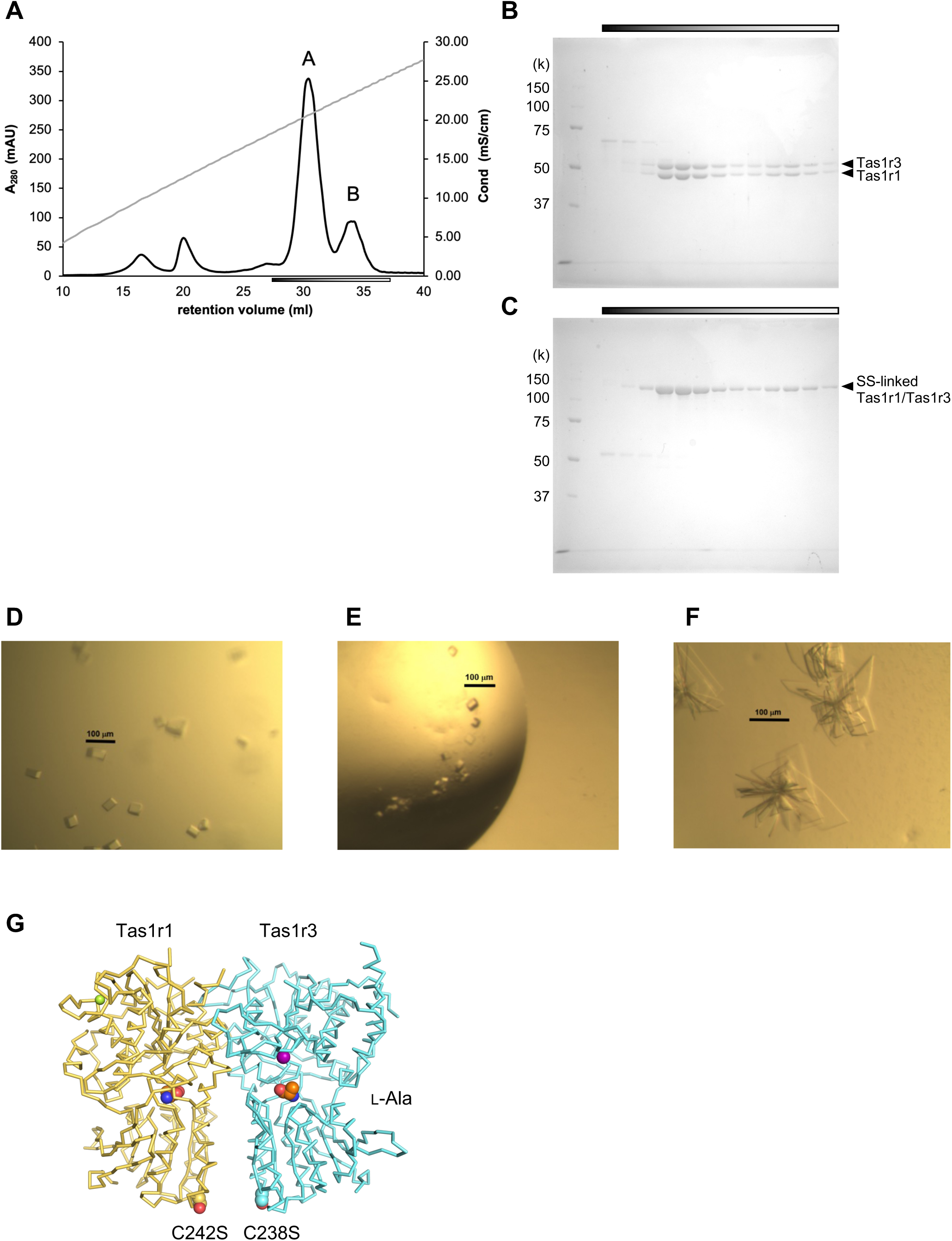
Sample preparation and crystallization of pufferfish Tas1r1/Tas1r3LBD. (A) Representative elution profile of pufferfish Tas1r1/Tas1r3LBD obtained by anion-exchange chromatography. (B, C) SDS-PAGE of pufferfish Tas1r1/Tas1r3LBD under reduced (B) and non-reduced (C) conditions. The bars under the elution profile in panel A and above the gels in panels B and C represent the range of fractions analyzed by SDS-PAGE. (D-F) Crystals of pufferfish T1r1/T1r3LBD in the L-alanine- (D), D- alanine- (E), and D-phenylalanine- (F) bound states. (G) Mutations introduced into the protein sample for crystallization to eliminate free cysteine residues at the surface.

**Extended Data Figure 2.**
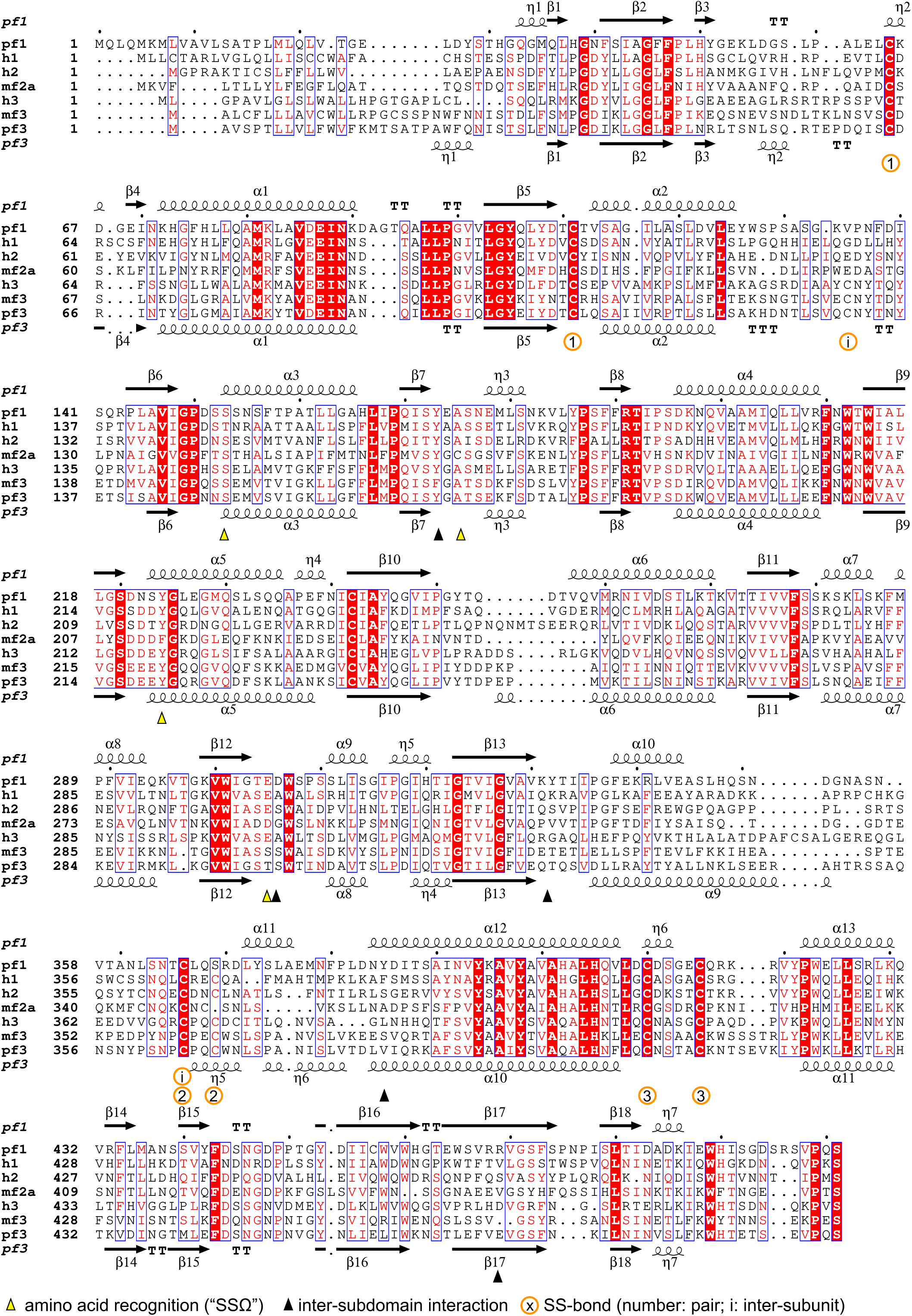
Amino acid sequence alignment of pufferfish Tas1r1 (pf1) and Tas1r3 (pf3); human TAS1R1 (h1), TAS1R2 (h2), and TAS1R3 (h3); medaka Tas1r2a (mf2a); and Tas1r3 (mf3).

**Extended Data Figure 3.**
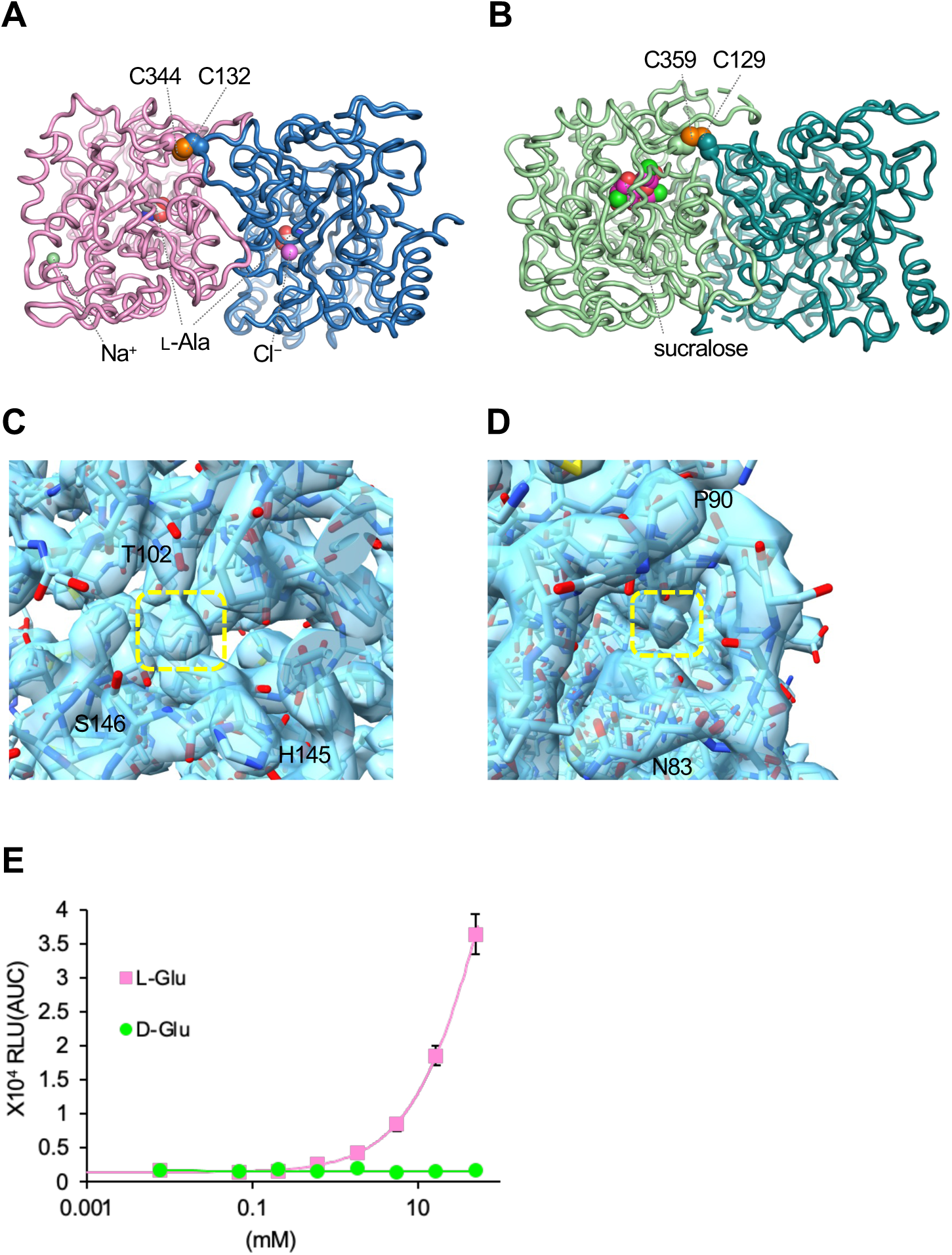
Structural and functional characteristics of the TAS1Rs. (A, B) Inter-subunit disulfide bonds in medaka Tas1r2a/Tas1r3LBD (A, PDB ID: 5X2N) and human TAS1R2/TAS1R3 (B; PDB ID: 9NOU). (C, D) Additional density blobs observed in the cryo-EM density map of human TAS1R2/TAS1R3 (EMDB: EMD-49167), demarcated in yellow dashed boxes. (C) Site corresponding to the chloride ion-binding site in TAS1R3. (D) Site corresponding to the sodium-ion binding site in TAS1R1. (E) Responses of full-length human TAS1R1/TAS1R3 evoked by L- and D-glutamate, analyzed under the same assay conditions shown in Fig. 2A in the main text. Values are the mean ± SEM at concentrations of 0.0076, 0.069, 0.21, 0.62, 1.9, 5.6, 17, and 50 mM from six experiments.

**Extended Data Figure 4.**
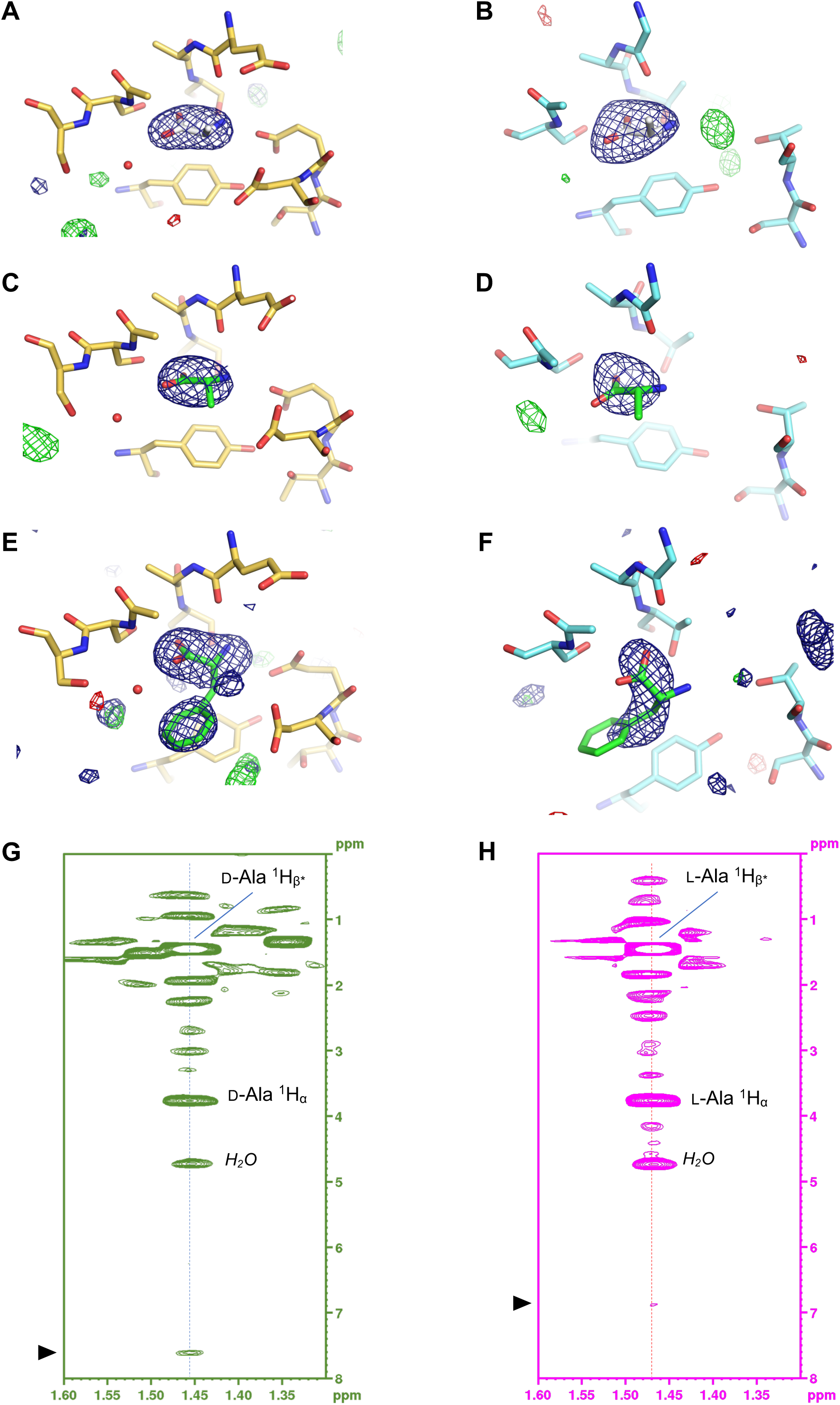
Amino acid-binding in pufferfish Tas1r1/Tas1r3LBD. (A–F) simulated annealing (SA)-omit electron density map and *F*_o_ – *F*_c_ difference Fourier map at the amino acid-binding sites. (A, B) L-Alanine-bound Tas1r1 (A) /Tas1r3 (B)-LBD structures with the SA-omit map at 4.0 *σ* and the *F*_o_ – *F*_c_ map at ± 3.5 *σ*. (C, D) D-Alanine-bound Tas1r1 (C) /Tas1r3 (D)-LBD structures with the SA-omit map at 3.2 *σ* and the *F*_o_ – *F*_c_ map at ± 3.5 *σ*. (E, F) D-Phenylalanine-bound Tas1r1 (E) /Tas1r3 (F)-LBD structures with the SA-omit map at 3.0 *σ* and the *F*_o_ – *F*_c_ map at ± 3.5 *σ*. (G, H) NMR spectra of D- and L-alanine-bound Tas1r1/Tas1r3LBD. ^13^C-edited-NOESY spectra of the Tas1r1/Tas1r3LBD – [2,3-^13^C_2_; ^15^N]-D-alanine (^13^C,^15^N-D-Ala) complex (G) and Tas1r1/Tas1r3LBD – [2,3-^13^C_2_; ^15^N]-L-alanine (^13^C,^15^N-L-Ala) complex (H) in the β-methyl proton region of D-/ L-alanine. NOE signals were observed along the dashed line in each spectrum, corresponding to protons spatially close to the methyl protons of ^13^C,^15^N- D/L-alanine (within approximately 5 Å). The intramolecular NOE signals of alanine and NOEs originating from the residual H_2_O were annotated in the spectra. Arrowheads indicate NOE signals, presumably between the methyl proton in D- or L-alanine and an aromatic ring.

**Extended Data Figure 5.**
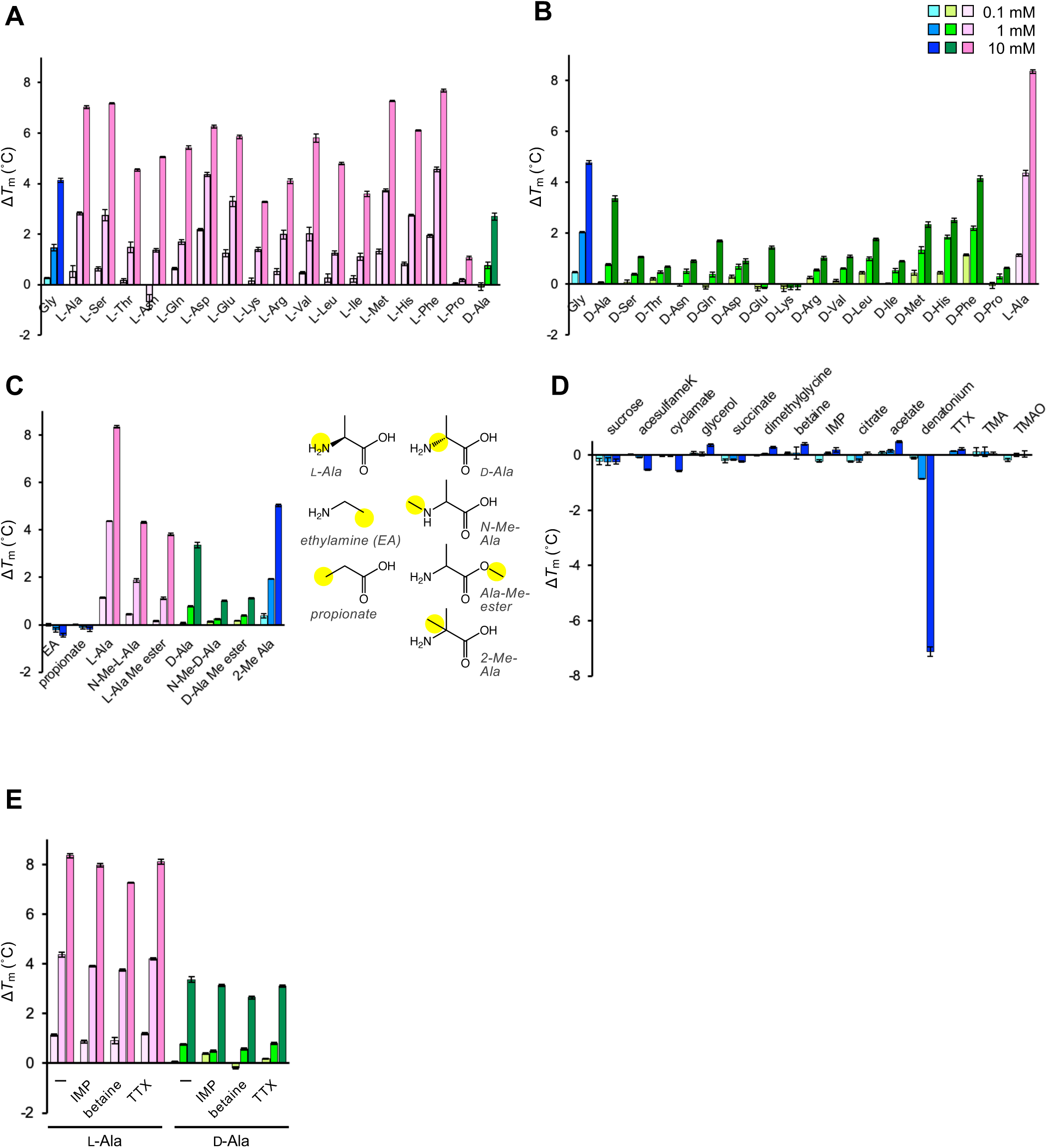
Ligand-binding analysis of pufferfish Tas1r1/Tas1r3LBD analyzed by DSF, in which binding was evaluated by an increase of melting temperature (Δ*T*_m_) of the protein ^23^. (A) l-amino acids; (B) d-amino acids; (C) alanine and their derivatives; (D) various chemicals; (E) alanine in the absence (–) and presence of inosine monophosphate (IMP), betaine, and tetrodotoxin (TTX), showing no enhancement of alanine binding.

**Extended Data Figure 6.**
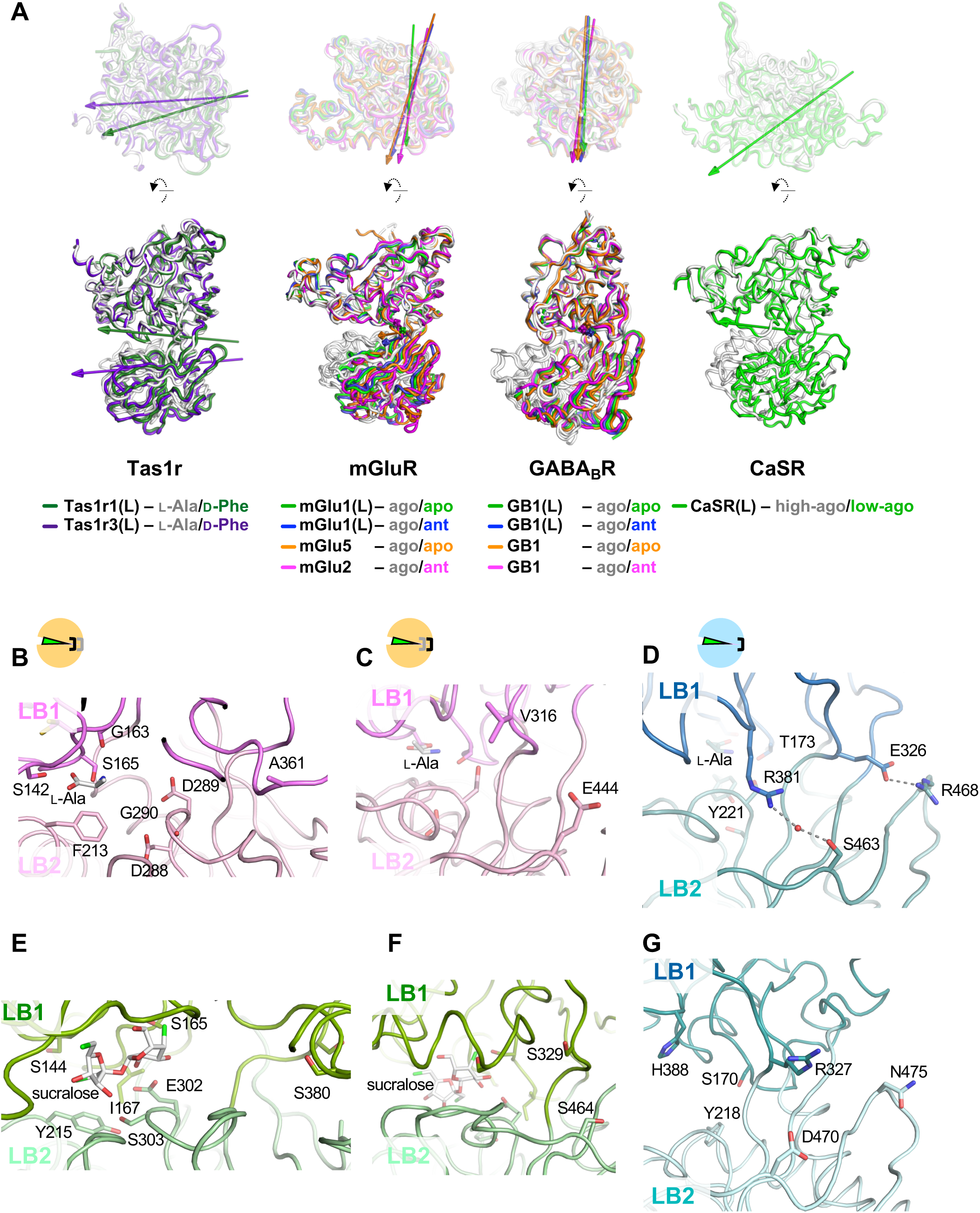
Domain motion and inter-subdomain interactions in LBDs of class C GPCRs. (A) Domain motion analysis of LBDs of class C GPCRs. Top and side views of the LBDs, with arrows representing the rotation axes of domain motions between two states with different ligands, are shown at the bottom of each panel. Domain motions were analyzed using DynDom ^63^. The coordinates used for the analysis are summarized in Extended Data Table 3. (B–G) Intersubdomain interactions in reported TAS1R structures. (B–D) Sites in the medaka Tas1r2a/Tas1r3 structure (PDB ID: 5X2N) corresponding to those shown in Fig. 4A (A; in Tas1r2a), 4 B (B; in Tas1r2a), and 4C (C; in Tas1r3). In Tas1r3, a water-mediated hydrogen bond between Arg381 (LB1) and Ser463 (LB2), and a salt bridge between Glu326 (LB1) and Arg468 (LB2) were observed. (E–G) Sites in the human TAS1R2/TAS1R3 structure (PDB ID: 9NOV) corresponding to those shown in Fig. 4A (D; in TAS1R2), 4 B (E; in TAS1R2), and 4C (F; in TAS1R3). Note that the TAS1R3 structure is in the “open” state, and no corresponding interactions were observed.

**Extended Data Table 1.**
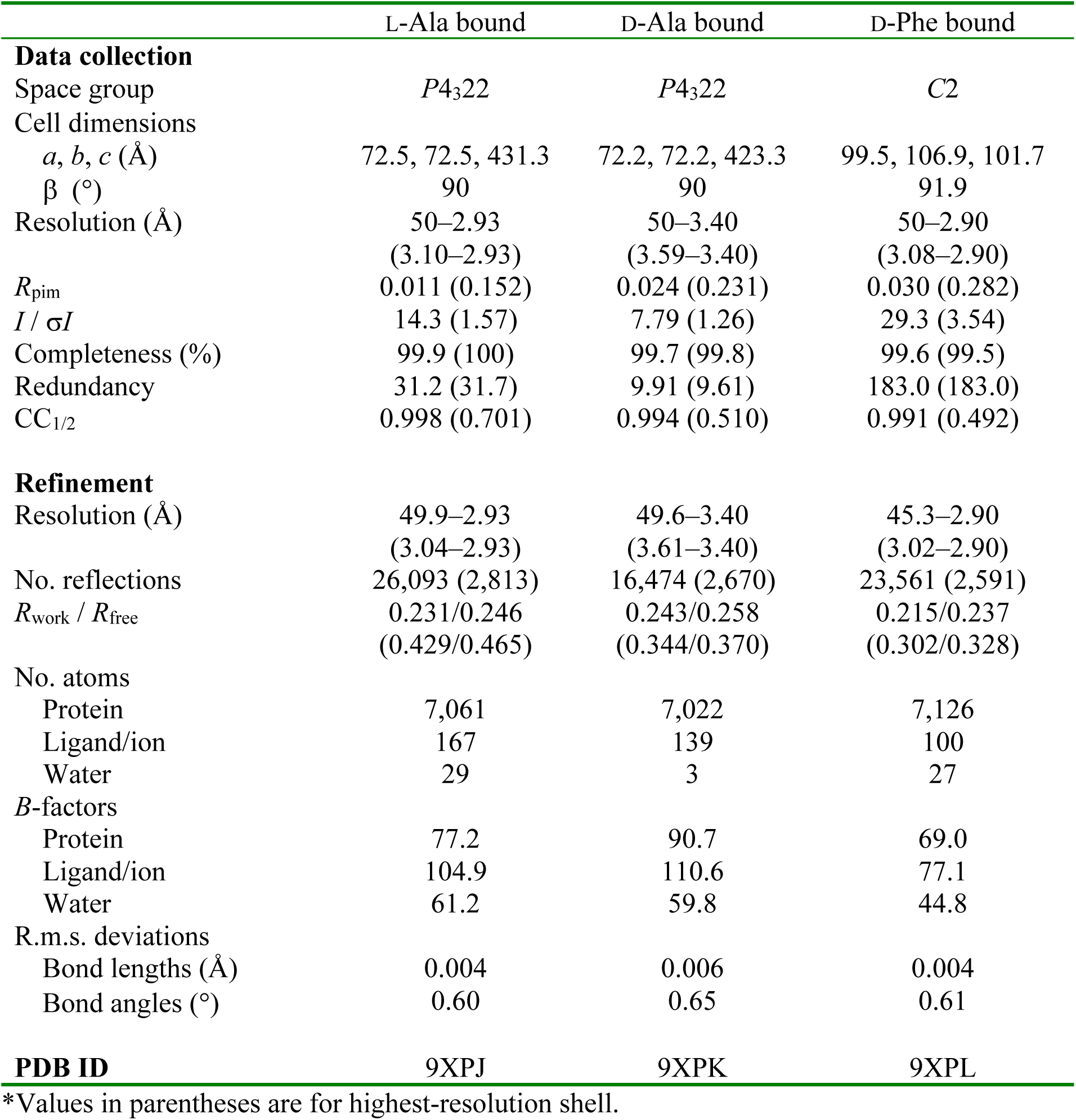
Data collection and refinement statistics.

**Extended Data Table 2.** Apparent dissociation constants of pufferfish Tas1r1/Tas1r3LBD determined by DSF.

| ligands | $K_{d-app}$ (mM) |
| --- | --- |
| L-alanine | $0.126 \pm 0.007$ |
| *N*-methyl-L-alanine | $0.345 \pm 0.008$ |
| L-alanine methyl ester | $0.867 \pm 0.158$ |
| D-alanine | $1.44 \pm 0.09$ |
| *N*-methyl-D-alanine | ND* |
| D-alanine methyl ester | ND* |
| 2-methylalanine | $1.29 \pm 0.14$ |

The values represent the mean ± SE of three independent experiments with three technical replicates. \*ND: not determined.

**Extended Data Table 3.** Conformational changes in the ligand-binding domains of class C GPCRs.

| Receptor* | Conformer 1 (closed) |  |  | Conformer 2 (open) |  |  | Conformation change |  |  |  |
| --- | --- | --- | --- | --- | --- | --- | --- | --- | --- | --- |
|  | PDB ID | Ligand<sup>†</sup> | Ligand type<sup>†</sup> | PDB ID | Ligand<sup>†</sup> | Ligand type<sup>†</sup> | Orientation of rotation axis<sup>‡</sup> (°) | Rotation angle<sup>§</sup> (°) | Translation<sup>§</sup> (Å) | Closure<sup>§</sup> (%) |
| mGlu1(L) | 1EWK | L-Glu | agonist | 1EWT | none | – | – | 21.6 | 0.6 | 97.8 |
| mGlu1(L) | 1EWK | L-Glu | agonist | 1ISS | S-MCPG | antagonist | 16.6 | 21.6 | 0.6 | 97.8 |
| mGlu5 | 6N51 | L-quisqualate | agonist | 6N52 | none | – | 15.7 | 30.8 | 0.8 | 99.9 |
| mGlu2 | 7MTS | L-Glu | agonist | 7MTQ | LY341495 | antagonist | 9.6 | 27.7 | 0.3 | 95.2 |
| GABA<sub>B</sub>R(L) | 4MS3 | GABA | agonist | 4MQE | none | – | 14.8 | 29.4 | –0.2 | 98.3 |
| GABA<sub>B</sub>R(L) | 4MS3 | GABA | agonist | 4MR7 | CGP54626 | antagonist | 14.8 | 27.5 | –0.4 | 98.7 |
| GABA<sub>B</sub>R | 6UO8 | SKF97541 | agonist | 6VJM | none | – | 18.2 | 32.1 | 0.0 | 94.2 |
| GABA<sub>B</sub>R | 7C7Q | baclofen | agonist | 7C7S | CGP54626 | antagonist | 21.3 | 26.8 | –0.4 | 96.7 |
| CaSR(L)<sup>¶</sup> | 5K5S | Ca<sup>2+</sup>, L-Trp | agonist | 5K5T | Low Ca<sup>2+</sup> | ND | 36.6 | 28.5 | –1.9 | 99.2 |
| **Tas1r1(L)<sup>§</sup>** |  | **L-Ala** | **agonist** |  | **D-Ala** | **agonist** | **ND** | **ND** | **ND** | **ND** |
| **Tas1r3(L)<sup>§</sup>** |  | **L-Ala** | **agonist** |  | **D-Ala** | **agonist** | **ND** | **ND** | **ND** | **ND** |
| **Tas1r1(L)<sup>§</sup>** |  | **L-Ala** | **agonist** |  | **D-Phe** | **agonist** | **60.3** | **9.3** | **–0.0** | **99.5** |
| **Tas1r3(L)<sup>§</sup>** |  | **L-Ala** | **agonist** |  | **D-Phe** | **agonist** | **73.1** | **17.8** | **–0.3** | **99.5** |

\*Receptor names with or without the suffix “(L)” indicate the structural information of the isolated extracellular domain analyzed by X-ray crystallography or those consisting of both extracellular and transmembrane domains analyzed by cryo-electron microscopy, respectively. <sup>†</sup>Those bound to ligand-binding domains are described. <sup>‡</sup>Relative orientation of the rotation axis for inter-subdomain cleft opening/closing to the reference vector (the rotation axis for the conformational change between agonist-bound (PDB ID: 1EWK) and apo (PDB ID: 1EWT) mGlu1. See the Methods section of the main text. <sup>§</sup>Results from DynDom analysis when the coordinates of ligand-binding domain portion A for each conformer were taken as inputs. In the case of homodimers, such as mGluRs and CaSR, chain A was used as a representative input. For GABA<sub>B</sub>R, subunit 1, which is responsible for agonist/antagonist binding, was used as the input. <sup>¶</sup>As of May 2024, the open conformation structure of the ligand-binding domain was solely observed in the crystallographic structure of the isolated extracellular domain, and no full-length cryo-EM structures with the open LBD conformation have been reported. Therefore, whether a similar conformational change is observed in the full-length receptor has not yet been verified. <sup>§</sup>This study. ND: Conformation changes were not detected.

**Extended Data Table 4.** Apparent dissociation constants of L- and D-alanine to pufferfish Tas1r1/Tas1r3LBD mutants determined by DSF.

| mutation | $K_{d-app}$ (mM)<sup>a</sup> | $n^b$ | Mean difference<sup>c</sup> | $P^d$ | 95% CI |
| --- | --- | --- | --- | --- | --- |
| L-alanine |  |  |  |  |  |
| wild type (WT)<sup>e</sup> | $0.126 \pm 0.007$ | 3 | — | — | — |
| E174A | $0.0614 \pm 0.0177$ | 5 | 0.065 | 0.0437 | 0.002 – 0.128 |
| D306A | $0.0979 \pm 0.0225$ | 3 | 0.028 | 0.657 | –0.042 – 0.099 |
| Y385F | $0.186 \pm 0.013$ | 5 | –0.061 | 0.0654 | –0.123 – 0.003 |
| R469V | $0.203 \pm 0.013$ | 5 | –0.076 | 0.0168 | –0.140 – –0.013 |
| D-alanine |  |  |  |  |  |
| wild type (WT)<sup>e</sup> | $1.44 \pm 0.09$ | 3 | — | — | — |
| E174A | $1.69 \pm 0.46$ | 5 | –0.26 | 0.978 | –1.87 – 1.36 |
| D306A | $3.14 \pm 0.91$ | 3 | –1.71 | 0.0662 | –3.51 – 0.10 |
| Y385F | $1.98 \pm 0.12$ | 5 | –0.54 | 0.773 | –2.15 – 1.07 |
| R469V | $1.88 \pm 0.14$ | 4 | –0.43 | 0.892 | –2.12 – 1.26 |

<sup>a</sup>The values are the mean ± SE. <sup>b</sup>Number of independent experiments with 2–4 technical replicates. <sup>c</sup>Mean differences in $K_{d-app}$ values between the WT and each mutant. <sup>d</sup> $P$ values from one-way ANOVA with Dunnett's test, with WT as the control group. <sup>e</sup>Data were derived from the same experiments shown in Extended Data Table 2.

**Supplementary Figure 1.**
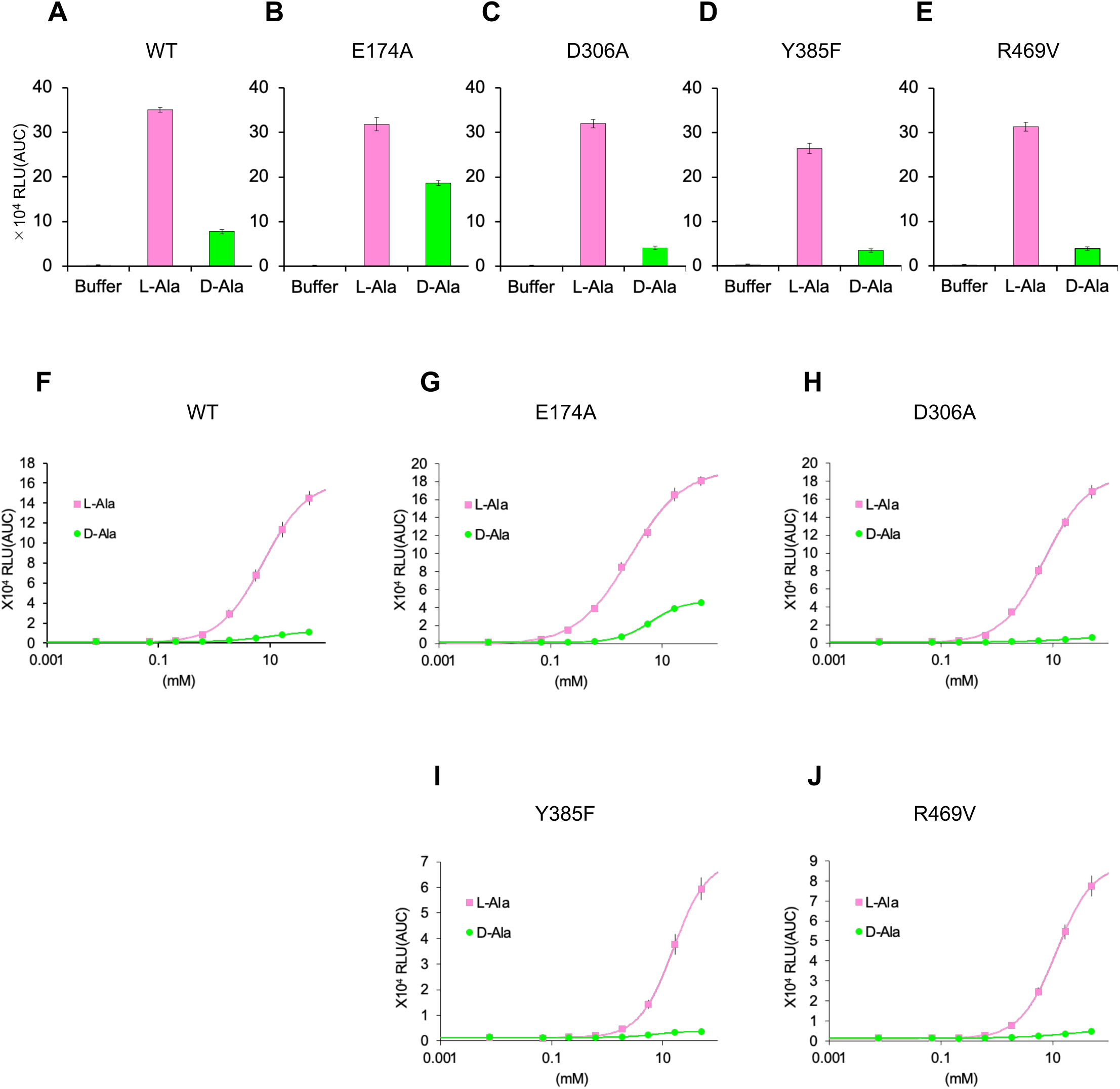
Responses to L- and D-alanine by pufferfish Tas1r1/Tas1r3 mutants. (A-E) Responses to L- and D-alanine at 50 mM by pufferfish Tas1r1/Tas1r3 mutants, used for Fig. 4F in the main text. Data are mean values ± SEM from six experiments. (H-J) Dose-response curves of L- and D-alanine by pufferfish Tas1r1/Tas1r3. Curves for wild type Tas1r1/Tas1r3 (A) and mutant receptors composed of Tas1r1-E174A/Tas1r3 (B), Tas1r1-D306A/Tas1r3 (C), Tas1r1-Y385F/Tas1r3 (D), and Tas1r1-R469V/Tas1r3 (E) are shown. Data are mean values ± SEM at the concentrations of 0.0076, 0.069, 0.21, 0.62, 1.9, 5.6, 17, and 50 mM from six experiments.

